# Intrinsic structure of model-derived metrics for *in silico* proarrhytmic risk assessment identified by global sensitivity analysis

**DOI:** 10.1101/543926

**Authors:** Jaimit Parikh, Paolo Di Achielle, James Kozloski, Viatcheslav Gurev

## Abstract

Multiscale computational models of heart are being extensively investigated for improved assessment of drug-induced Torsades de Pointes (TdP) risk, a fatal side effect of many drugs. Model-derived metrics (features) such as action potential duration and net charge carried by ionic currents (*qNet*) have been proposed as potential candidates for TdP risk stratification after being tested on small datasets. Unlike purely statistical approaches, model-derived metrics are thought to provide mechanism-based classification. In particular, the underlying mechanism behind the success of the recently proposed *qNet* metric is attributed to its correlation to early afterdepolarizations (EADs), which are known to be cellular triggers of TdP. Analysis of critical model components and of ion-channels that have major impact on model-derived metrics can lead to improvement in the confidence of the prediction. In this paper, we analyze a large population of virtual drugs to systematically examine the influence of different ion channels on model-derived metrics that have been proposed for proarrhythmic risk assessment. Global sensitivity analysis (GSA) methods were employed to determine and highlight the critical input parameters that affect different model-derived metrics. We observed significant differences between the sets of input parameters that control model-derived metrics and generation of EADs in the model, thus opposing the idea that these metrics and sensitivity to EAD might be strongly correlated. Moreover, in classification of a small set of actual drugs, we found that the classifiers based on EADs performed worse than those built on other model-derived metrics. Hence, our analysis points towards a need for a better mechanistic interpretation of promising metrics such as *qNet* based on formal analyses of models. In particular, GSA should constitute an essential component in the *in silico* workflow for proarrhythmic risk assessment to yield improved understanding of the structure of mechanistic dependencies surrounding model-derived metrics while ultimately providing increased confidence in model-predicted risk.

## 1 INTRODUCTION

Drug-induced Torsades de Pointes (TdP) is a specific form of polymorphic ventricular tachycardia that leads to ventricular fibrillation and sudden cardiac death (Yap and Camm, 2003). Several drugs have been withdrawn from the market in the past due to TdP risk (Gintant, 2008). Although the current clinical safety guidelines are successfully preventing drugs with torsadogenic risk from reaching the market (Sager et al., 2014), safe drugs may be potentially excluded due to the low specificity of the screening process, which targets only hERG channels. The Comprehensive *in vitro* Proarrhythmia Assay (CiPA) is a global initiative to provide revised guidelines for better evaluation of the proarrhythmic risk of drugs (Fermini et al., 2016). *In silico* evaluation of proarrhythmic action for different compounds constitutes an important foundation under the CiPA initiative to link data from *in vitro* assays to changes in cell behavior (Fermini et al., 2016; Colatsky et al., 2016).

The main component of the *in silico* evaluation are classifiers that are based on the so-called “derived features”, input variables for the classifiers that are extracted from the outputs of biophysical models. The term “direct features” refers instead to the original feature set estimated from experiments investigating how drugs affect ion channel kinetics. Biophysical models serve as complex transformations that generate feature sets conditioned to the prior knowledge used in creating the model, thus potentially improving the efficacy of linear classifiers in inferring TdP risk. Diverse sets of derived features have been suggested as potential candidates for TdP risk classification (Table 1). In one of the earliest works on the use of the myocyte models for TdP risk prediction, simulated action potential duration at 90 % repolarization (*APD*90) was shown to provide the best discrimination of torsadogenic and non-torsadogenic drugs (Mirams et al., 2011). Other derived features extracted from the action potential (e.g., early after depolarization (EAD) and transmural dispersion of repolarization (TDR)), have also been suggested as possible candidate metrics for TdP risk prediction (Christophe, 2013, 2015). Considering derived features from calcium transient in addition to features of the action potential have been shown to improve TdP risk discrimination (Lancaster and Sobie, 2016). Recently, tertiary TdP risk classifiers trained on a set of 12 drugs categorized into 3 clinical TdP risk groups (high, intermediate, and low/no risk) have been developed at FDA as a part of the CiPA initiative (Li et al., 2017; Dutta et al., 2017). Finally, two new derived features *cqInward* (Li et al., 2017) and *qNet* (Dutta et al., 2017) have been proposed to separate the 12 training drugs into desired target groups. The *qNet* metric was further validated on 16 test compounds (Li et al., 2018). Uncertainty quantification methods (Johnstone et al., 2016) have recently gained increased attention due to their ability to better estimate the confidence of the model-predicted risk (Chang et al., 2017) by taking into account noise in the *in vitro* measurements of drug-induced effects on ionic currents, under the CiPA initiative.

**Table 1.**
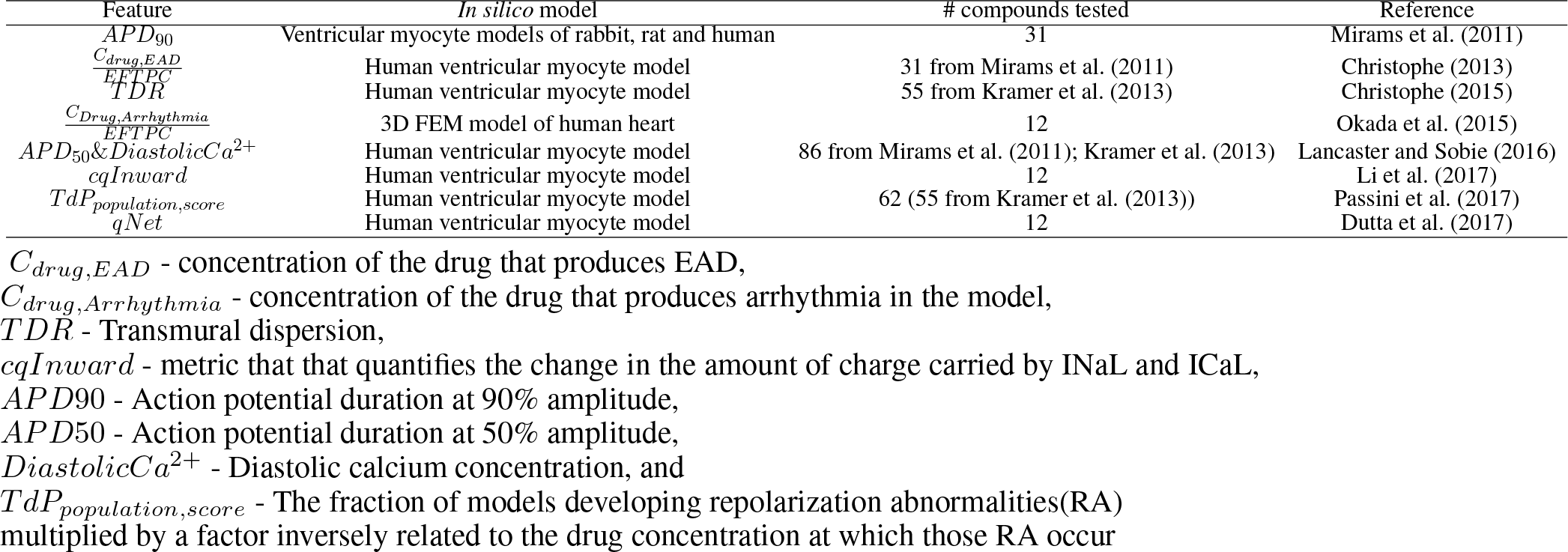
Previously proposed derived features.

Model-derived features that are directly linked to drug-induced changes in myocyte membrane activity are thought to provide mechanism-based classification of compounds into different risk categories by providing possible insights into TdP mechanisms. The *qNet* metric is thought to provide a measure of propensity of myocytes to undergo EADs (Dutta et al., 2017; Chang et al., 2017), that are known to be the trigger of TdP (Yan et al., 2001). In this paper, we apply global sensitivity analysis (GSA) to the existing CiPA *in silico* framework to identify key model components that require special treatment for reducing uncertainties in the estimated model-derived metrics and, ultimately, TdP risk classification. Unlike previous approaches where the initial feature selection and construction were performed by testing on a small set of drugs, in this study, we analyzed a large virtual population of drugs to identify the critical input parameters regulating the variation of several previously proposed model-derived metrics (e.g., *APD*90, *qNet*) for proarrhythmic risk classification. We also compare the key inputs that regulate these model-derived metrics to those regulating generation of the EADs. We demonstrate that, in spite of previously claimed ties between *qNet* and EAD generation, the parameters that affect *qNet* are different than those influencing cell sensitivity to EAD. Moreover, we show that classifiers built on EAD metrics perform worse than classifiers built on *qNet*. Hence, our results highlight the need for better mechanistic understanding of promising model-derived metrics. Furthermore, the sensitivity analysis results provide an explanation of the equivalent performance of direct and derived features.

## 2 METHODS

The CiPAORd model and input parameters section describes the *in silico* model used in the paper. To perform GSA, we generated a large set of virtual drugs. A virtual drug comprises a random vector of changes to parameters of ion channels of the model. The details of the input parameters considered for generating the virtual drug population are presented in Sampling virtual drug population. Responses to the virtual drugs were examined, evaluating several model-derived features such as *APD*_90_, *qNet*, and peak calcium concentration (*peakCa*). The section *In silico* simulations and derived features presents details on the derived features extracted from the *in silico* model. To explore the link between model-derived metrics and EADs in the model virtual drugs were also tested for their ability to induce EAD. In the section EAD protocols we discuss the protocols used to test for EAD generation in the model. The methods used for GSA are described in the Global sensitivity analysis. Finally, the methods for classifying the 28 drugs selected under the CiPA initiative, which we refer in the manuscript as “CiPA drugs”, with respect to their defined torsadogenic risk are described in the section Tertiary risk stratification of “CiPA drugs”.

### CiPAORd model and input parameters

In this study, we perform GSA on the CiPAORd model (Dutta et al., 2017). The CiPAORd model was developed at FDA by introducing several modifications to the original O’Hara-Rudy ventricular myocyte model (O’Hara et al., 2011) to improve proarrhythmic risk assessment.

Several input parameters have been used for simulation of virtual drug effects. For the hERG channels, we used the concentration response of the drug, *E*_*max*_, the unbinding reaction rate, *K*_*u*_, and the membrane voltage at which half of drug-bound channels are open, *V*_*half*_, as input parameters for the model. In this paper, we refer to the *E*_*max*_ parameter that represents the static component of the hERG block as *sbIKr*. For the other channel currents (i.e., fast sodium current *INa*, late sodium current *INaL*, L-type calcium channel current *ICaL*, slow-rectifying potassium channel current *IKs*, inward rectifier potassium current *IK*1, transient-outward current *Ito*) we used the general Hill equation of channel block,

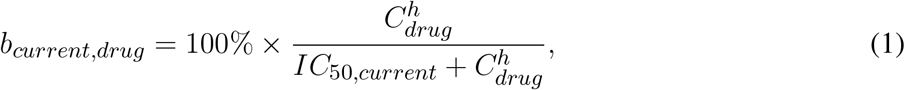

where current = {*INa*, *INaL*, *ICaL*, *IKs*, *IK*1, *Ito*}, *IC*_50,*current*_ is the drug concentration at which a current is reduced by half, *C*_*drug*_ is the drug concentration, and *h* is the Hill coefficient. The drug-induced blocks of channel currents *b*_*current,drug*_ are used to scale the maximum conductance of the current *g*_*current*_ in the *in silico* model calculated as

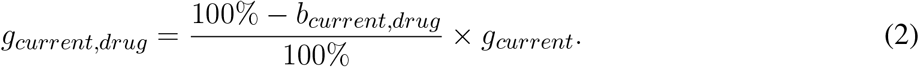

We perform GSA explicitly with respect to *b_current,drug_* rather than *IC*_50*,current*_, *C*_*drug*_, and *h*. In this study, we refer to the parameters of the block of *INa*, *INaL*, *ICaL*, *IKs*, *IK*1 and *Ito* as *bINa*, *bINaL*, *bICaL*, *bIKs*, *bIK*1, and *bIto*, respectively. Equation (1) is used in classification of real compounds.

### Sampling virtual drug population

The population of virtual drugs is created through Monte Carlo sampling from a high-dimensional (10-D) input parametric space. The parametric space represent changes in model parameters used to describe drug binding and blocks of ionic currents. Basic cycle length (BCL) of cell pacing in the simulations was also considered as a parameter for GSA. The input parameters and their examined ranges is provided in (Table 2). In some cases, GSA was performed on metrics derived from outputs of a mid-myocardial cell (defined as M cell in O’Hara et al. (2011)) model. M cells are very sensitive to blocks of repolarization currents and produce EAD more easily. Sensitivity to EADs makes the analysis more complicated, and the range of hERG block for M cells had to be accordingly reduced.

**Table 2.**
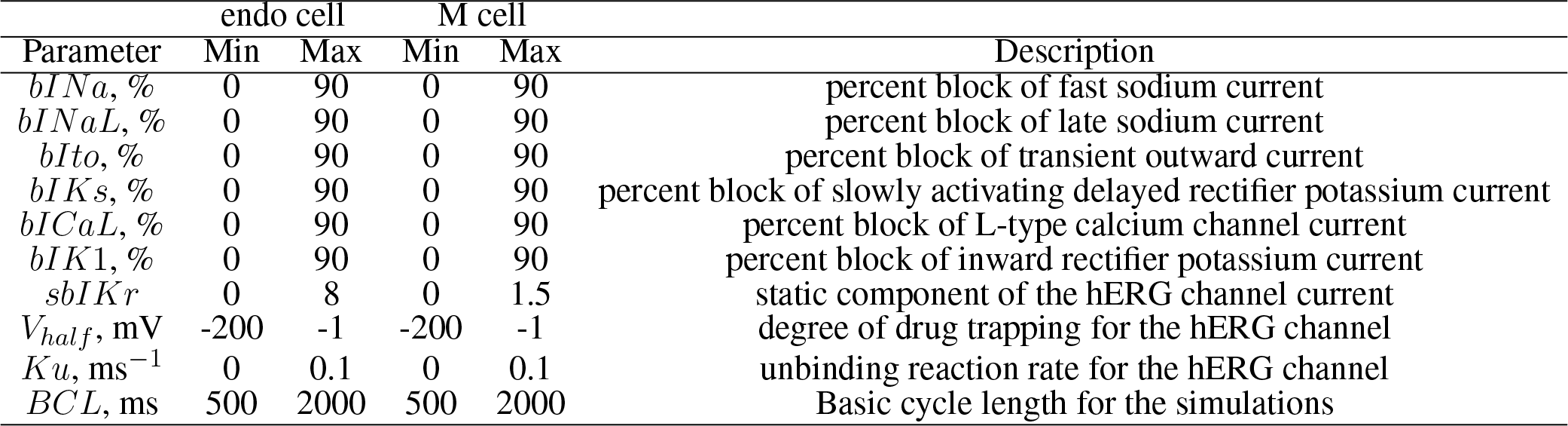
Ranges of input parameters

### *In silico* simulations and derived features

The action potential and calcium transients of the cells were simulated for the large virtual population of drug dataset (*>*20000 drugs) generated for GSA, and, separately, for the CiPA training (12 drugs) and validation (16 drugs) datasets (manual patch clamp data) (Li et al., 2017; Dutta et al., 2017) using the CiPAORd model. Model simulations were run for 200 beats to achieve pseudo steady state. Simulations were carried out at different pacing rates (a parameter in GSA) for each of the endocardial (endo), mid-myocardial (M), and epicardial (epi) cell types. Several standard metrics explored previously for TdP risk discrimination were calculated from the action potential and *Ca*^2+^ transients. The metrics obtained from the *in silico* models are listed in the Table 3. Note that the metric *qNet* was calculated as the area under the curve traced by the net current (*Inet* = *ICaL* + *INaL* + *IKr* + *IKs* + *IK*1 + *Ito*) from the beginning to the end of the last simulated beat as defined in Dutta et al. (2017).

**Table 3.** Derived features extracted from the CiPAORd model

| Derived Feature | Description |
| --- | --- |
| $qNet$ | Net electronic charge carried by $IKr$, $INaL$, $ICaL$, $IKs$, $IK1$, $Ito$ currents |
| $APD_{90}$ | Action potential duration at 90% repolarization |
| $APD_{50}$ | Action potential duration at 50% repolarization |
| $peakVm$ | Peak voltage |
| $diastolicCa$ | Diastolic calcium level |
| $peakCa$ | Peak value of intracellular calcium |
| $CaTD_{50}$ | Calcium transient duration at 50% return to baseline |
| $CaTD_{90}$ | Calcium transient duration at 90% return to baseline |

### EAD protocols

Drug-induced EAD risk (sensitivity of a cell against EAD generation) for both the virtual drugs and the CiPA compounds was examined in the endo and M cell types using two separate protocols. The M cell type in the CiPAORd model was more prone to EAD generation than the endo cell type. We tested generation of pause-induced EADs (that are implicated as triggers of TdP (Yan et al., 2001; Liu and Laurita, 2005; Viswanathan and Rudy, 1999)) in the M cell type as in our previous study (Parikh et al., 2017). Briefly, the cell was stimulated 200 times at a constant cycle length. After 200 stimuli, an additional stimulus was applied following a pause equal to the basic cycle length. In the endo cells, pause-induced EADs occurred rarely, and we examined drug-induced EAD risk in presence of an added perturbation by reducing the maximum conductance of hERG channel current (*IKr*) as in (Dutta et al., 2017). The cell was stimulated for 200 beats with additional block of maximum conductance of *IKr* by 85%. The 85% block was selected since almost half of the population of virtual drugs (across the entire parametric space observed) resulted in EAD development in the model simulations.

### Global sensitivity analysis

GSA was performed using a variance-based sensitivity method (Saltelli et al., 2008; Sobol’, 2001)), and Monte Carlo filtering Hornberger and Spear (1981); Saltelli et al. (2008). The supplemental material also reports analysis using Morris methods for comparision.

#### Variance-based global sensitivity analysis

Sobol sensitivity analysis method (Sobol’, 2001) is a model-independent GSA method that is based on variance decomposition. It relies on an all-at-time (AAT) sampling strategy where output variations are induced by varying all the input factors simultaneously. Let a derived-metric *Y* from a computational model be represented by a function *f* of the input parameters,

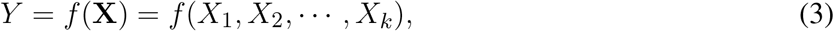

where **X** = {*X*_1_*, X*_2_ *X*_*k*_} is the input parameter set. The function can then be decomposed into a sum of elementary functions of increasing dimensions,

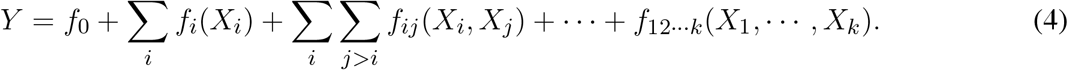

The input parameters are assumed to be random variables that are uncorrelated and mutually independent. The functional decomposition can be translated into a variance decomposition. This allows to quantify the variance contribution to the total output of individual parameters and the parameter interactions,

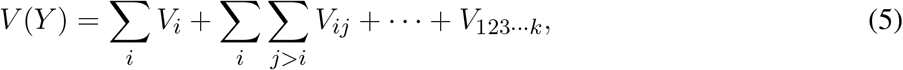

where *V*_*i*_ = *V*_*X*_*i*__ [*E*_*X*_*∼i*__ (*Y*|*X*_*i*_)] is the first-order effect for a given model input *X*_*i*_, *V*_*ij*_ = *V*_*X*_*i*__,*X*_*j*_ [*E*_*X*_*∼ij*__ (*Y*|*X*_*i*_, *X*_*j*_)] − *V*_*X*_*i*__ [*E*_*X*_*∼i*__ (*Y*|*X*_*i*_)] − *V*_*X*_*j*__ [*E*_*X*_*∼j*__ (*Y*|*X*_*j*_)] and so on are the higher-order effects due to interactions of model inputs. Here, *E*_*X*_*i*__, *V*_*X*_*i*__ are expectation and variance taken over *X*_*i*_; *X*_*∼i*_ denotes all factors but *X*_*i*_. The Sobol sensitivity indices are obtained as the ratio of partial variance to the total output variance,

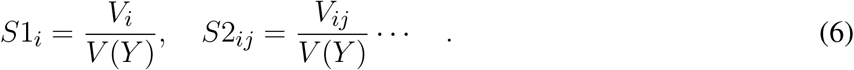

The number of sensitivity indices in (6) grow exponentially with *k* and typically only sensitivity indices of up to order two (*S*1_*i*_and *S*2_*i*_) and the total-effect indices (*ST*_*i*_) are estimated (Iooss and Lemaître, 2014). The total-effect index

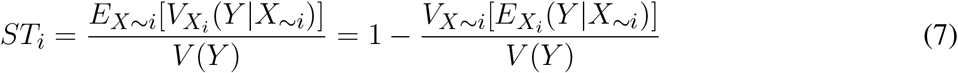

measures the impact of main effect of *X*_*i*_ and all its higher-order interaction effects with the other parameters (Homma and Saltelli, 1996). The Python SALib package was employed to perform the variance-based sensitivity analysis (Herman and Usher, 2017). The calculations of *S*1_*i*_, *ST*_*i*_ and *S*2_*ij*_require *n* × (2*k* + 2) model evaluations using Saltelli’s sampling scheme (Saltelli, 2002) where *n* is the sample size and *k* is the number of input parameters. In this study, we considered *n* = 1000 unless otherwise specified, resulting in 22000 Monte Carlo samples (virtual drugs) for *k* = 10.

Multivariate linear regression has been used in the past (Sobie, 2009) to identify sensitivity of outputs from cardiac cell models to changes in input parameters. To illustrate the differences between linear regression ^1^ and variance-based sensitivity analysis, in the Figure 1, we provide few examples highlighting the differences between the variance based sensitivity measures and sensitivity coefficients from the linear regression. For a hypothetical output feature (Feature1 in Figure 1 A) that can be perfectly fitted by a linear regression of model input parameters (*Feature*1 = 1.5*P*_1_ + *P*_2_ + 5) the sensitivity coefficients obtained using the two methods are identical (Figure 1). In contrast, the sensitivity estimates are inaccurate for the model features that present non-linear input-output relationship when using the linear regression methods, and the variance-based analysis provides a proper estimate under this situation. The metrics *S*1 captures the contribution of the first order as well as all higher order terms of individual input. For the *Feature*2 in the Figure 1 the *S*1 terms capture the contribution of the *P*_1_ and 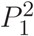 terms. The metric *S*2 captures all the second order interaction terms (i.e., *P*_1_*P*_2_). The variance in the hypothetical *Feature*3 in the Figure 1 depends on interaction between *P* 1 and *P* 2 parameters, which is captured by the *S*2 index, and also in the total sensitivity index *ST*, which includes all higher-order interaction terms, including *S*2. The *S*2 index of 0.38 indicates a contribution of 38% in the variance of *Feature*3 from the second-order interaction term (Figure 1 B). Hence, the variance based sensitivity analysis provides a method, which allowed us to estimate the contributions of parameter interactions and non-linear effects on regulation of the output features.

**Figure 1.**
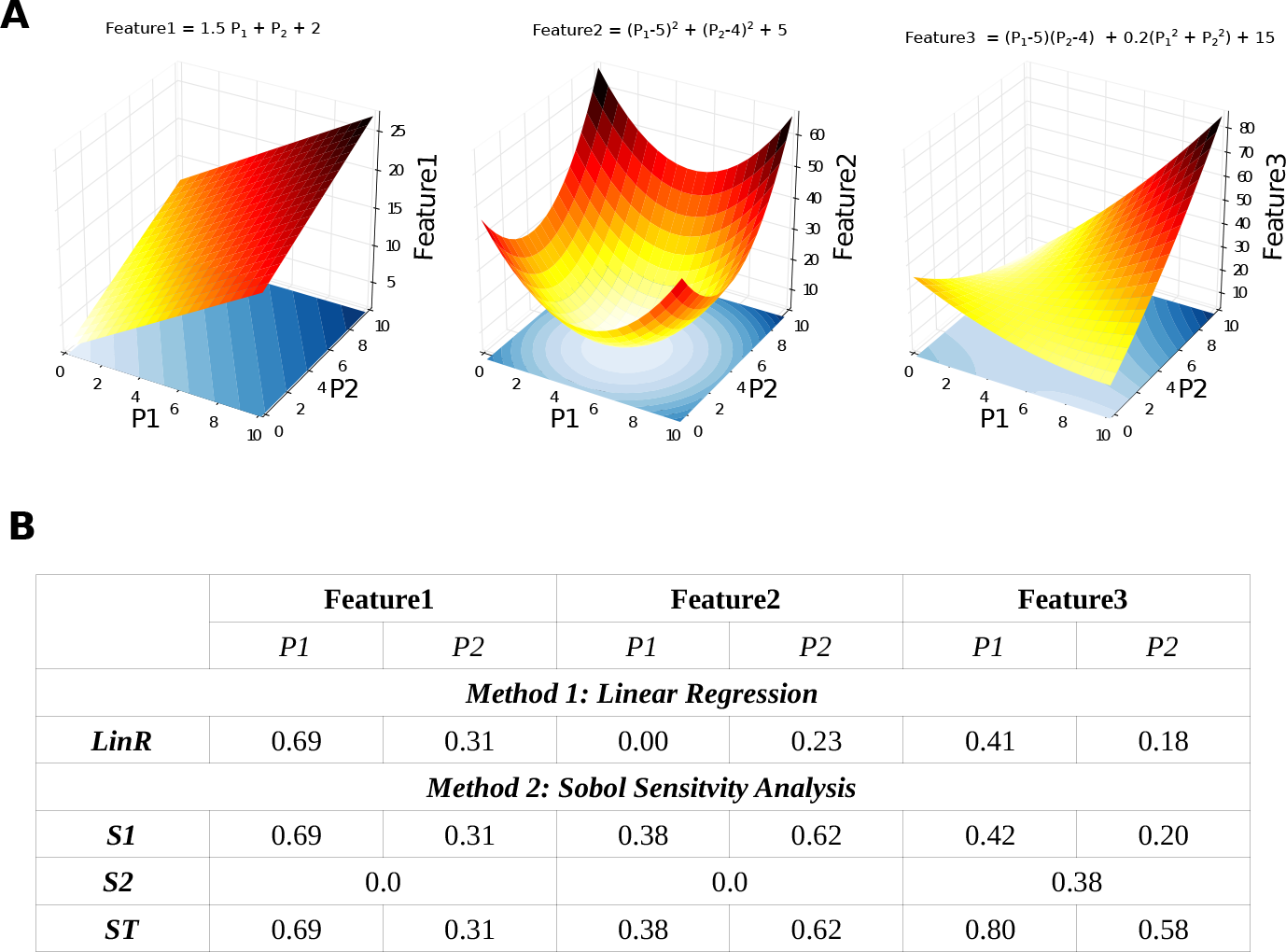
Example highlighting the difference between the multivariate linear regression and variance-based sensitivity methods. **A:** Schematic of variation in three synthetic features due to variation in two input parameters. **B:** Sensitivity estimates of the three synthetic features from **A** using multivariate linear regression and variance-based sensitivity methods.

#### Monte Carlo filtering

Monte Carlo filtering (MCF) is used generally in factor-mapping tasks to identify key input parameters responsible for driving model outputs within or outside predefined target regions (refer to (Saltelli et al., 2008) for a detailed methodology). A brief overview of the MCF technique in the context of EAD sensitivity analysis of the CiPAORd model is presented here. Model simulations were carried out for *n* Monte Carlo samples (virtual drugs) generated for the Sobol sensitivity analysis in presence of additional perturbations (see section EAD protocols). Each input parameter *X*_*i*_ of the Monte Carlo input sample set with size *n* is categorized into the “Behavioral” subset (*X*_*i*_|*EAD−*) and the “Non-behavioral” (*X*_*i*_|*EAD*+) with sizes *n*_1_ and *n*_2_ = *n − n*_1_, respectively, based on the absence and presence of EADs in simulated output. Empirical cumulative density functions (CDF) *F*_*n*_1__(*X*_*i*_|EAD−) and *F*_*n*_2__(*X*_*i*_|*EAD*+) are estimated for both the subsets of random input samples. The distance between the two empirical CDFs provides an estimate of sensitivity of *EAD* feature to the input parameter *X*_*i*_. Kolmogorov-Smirnov two-sample statistic test was used to quantify the difference between the two CDFs. Kolmogorov-Smirnov test is characterized by a D-statistic and a p-value. The D-statistic is defined as (Saltelli et al., 2008)

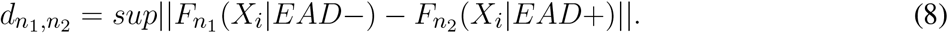

The larger the D-statistic (or equivalently the smaller the p-value), the more important the input parameter is in driving the behavior of the model to EAD (Saltelli et al., 2008). The sensitivity of *EADs* to different input parameters has been recently analyzed using multivariate logistic regression (Morotti and Grandi, 2016). Unlike logistic regression that relies on underlying assumption that a hyperplane separates the regions of interest, the MCF methods are valid in the more general case, where a highly non-linear or discontinuous surface separates the regions of interest (see Supplemental Material for comparision of the Monte Carlo filtering and logistic regression methods).

### Tertiary risk stratification of “CiPA drugs”

*In silico* simulations of blocks with the 28 “CiPA drugs” were carried out using the *in vivo* manual patch clamp measurements collected on the pharmacological effects of these compounds reported in Li et al. (2017, 2018). The effective therapeutic concentrations, the *IC*_50_ values, the Hill coefficient values, the drug binding parameters, and the defined torsadogenic risk of the “CiPA drugs” are listed in the Supplemental Material.

The “CiPA drugs” were classified based on the concentration (normalized to effective free therapeutic plasma concentration (EFTPC)) necessary to induce EADs and in the CiPAORd model. The effects of “CiPA drugs” were simulated using protocols described in the EAD protocols section at progressively increasing drug concentrations until EADs were observed. A maximum concentration of 70xEFTPC was tested for each drug. Drugs that did not result in EADs in the tested concentration range were classified as low risk drugs. Drugs that instead resulted in EADs at concentrations smaller than 8xEFTPC were labeled as high risk, while the remaining drugs were labeled as intermediate risk drugs. The threshold of 8xEFTPC was chosen to give best fit to the data. The “CiPA drugs” were also classified based on additional hERG channel perturbations that are required to induced EADs in the CiPAORd endo cell model as in Dutta et al. (2017). “CiPA drugs” were simulated using protocols described in the EAD protocols section at progressively increasing hERG channel perturbations (65 −100% block). Drugs that did not result in EADs in the presence of additional hERG channel perturbations were classified as low risk drugs. Drugs that instead led to EADs at perturbation levels smaller than or equal to 90% additional hERG block were labeled as high risk, while the remaining drugs that resulted in EADs in the presence of additional hERG block of *>*90 - 100% were labeled as intermediate risk drugs to achieve the best risk stratification. The classification of the “CiPA drugs” based on the *qNet* metric was also performed for comparison at 2xEFTPC concentration. The threshold values of 57 and 74, which provided the best discrimination across the different risk categories were used to classify the drugs into low, intermediate and high risk groups. Drugs with *qNet* values less than 57 were classified as high risk and drugs with *qNet* values greater than 74 were classified as low risk drugs.

## 3 RESULTS

### Analysis of global sensitivity for *APD*90, *qNet*, *peakCa*, and *EAD*

#### Variance-based analysis

Figure 2 demonstrates distribution of one of the model-derived metrics obtained from simulation of 22000 virtual drugs. The *APD*90 metric values are plotted against individual input parameters to visualize the influence of different input parameters on the metric. Each point on the scatter plot represents an individual virtual drug. Virtual drugs with comparable block of a particular ion-channel can result in a completely different output response due to differences in the effect of a virtual drug on other input parameters. The latter appears on the scatter plot as the variability of *APD*90 along the Y-axis. The scatter plot (Figure 2) shows a clear trend in *APD*90 with increase in the *sbIKr* parameter. This observed trend suggests that the parameter *sbIKr* is highly influential in regulating *APD*90. The Sobol sensitivity indices quantify the influence of individual parameters on the derived metrics.

**Figure 2.**
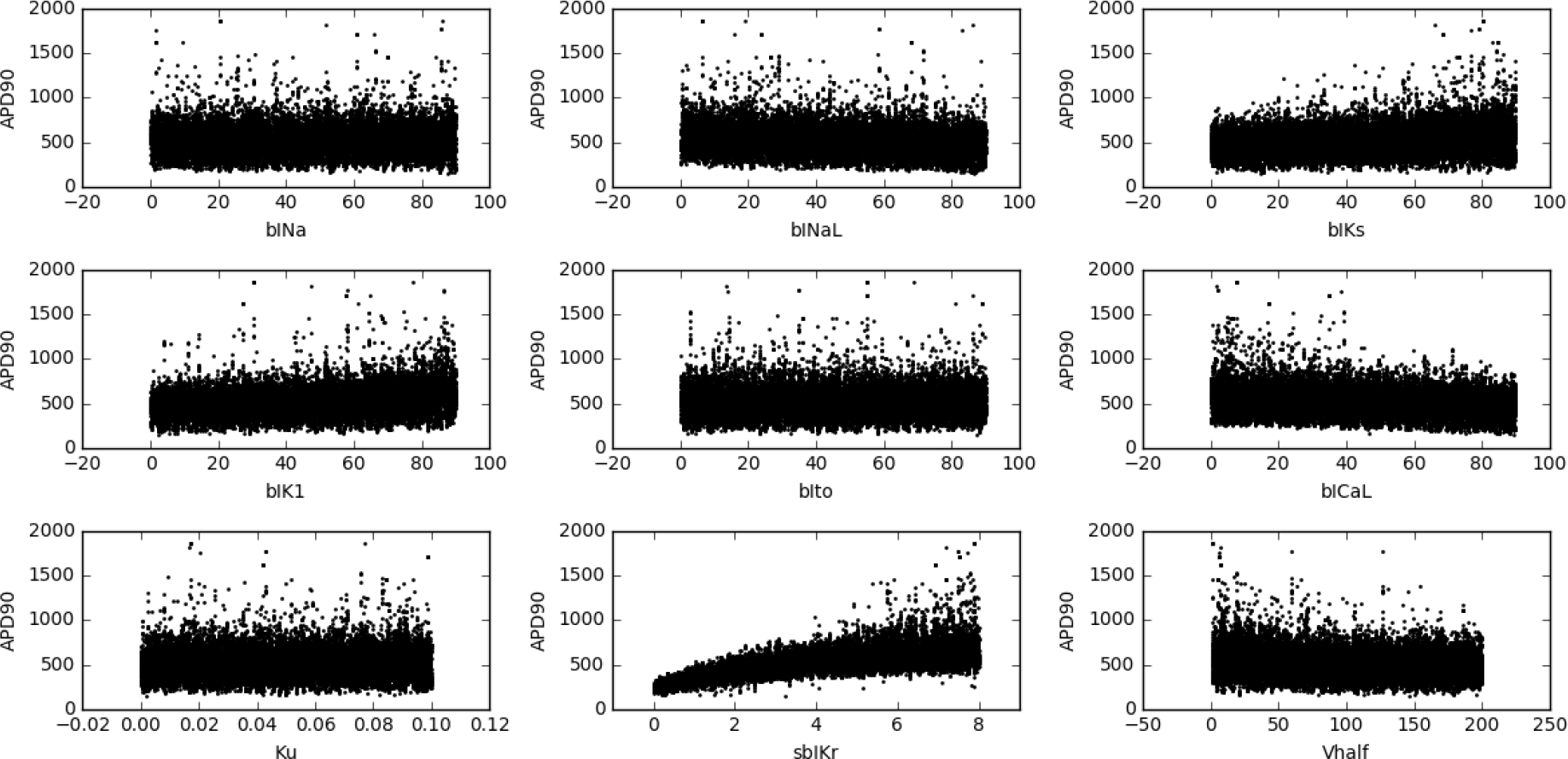
Scatter plot of *APD*90 versus different input parameters (direct features) for the 22000 simulated virtual drugs (endo cell model).

Figure 3A shows values of the first-order Sobol sensitivity indices (*S*1) and differences between total sensitivity indices (*ST*) and *S*1 for three output responses *APD*90, *qNet*, and *peakCa* simulated in the CiPAORd endo cell model. The Sobol sensitivity indices indicate that *APD*90 is the most sensitive to *sbIKr* block, *qNet* to *sbINaL*, and *peakCa* to *bICaL*. Sobol indices quantify the contribution of the input parameter to the metrics in isolation as well as in presence of interaction with other parameters. The effect of *sbIKr* on *APD*90 as quantified by *S*1 indicates that *sbIKr* contributes to *>* 50% of the variation observed in *APD*90 across the observed input space. *qNet* was found to be most sensitive to *bINaL*, *sbIKr*, *bINa*, and *bICaL* with contributions to the output variation of 30%, 26%, 17%, and 10%, respectively. *bICaL* had the strongest impact on the variability of *peakCa* concentrations with an *S*1 index of around 0.6. Among the different drug-effects evaluated via *in vitro* ion-channel screening, the changes in the block of transient outward current and dynamic hERG kinetic parameters showed relatively minor influence on the tested model-derived metrics.

**Figure 3.**
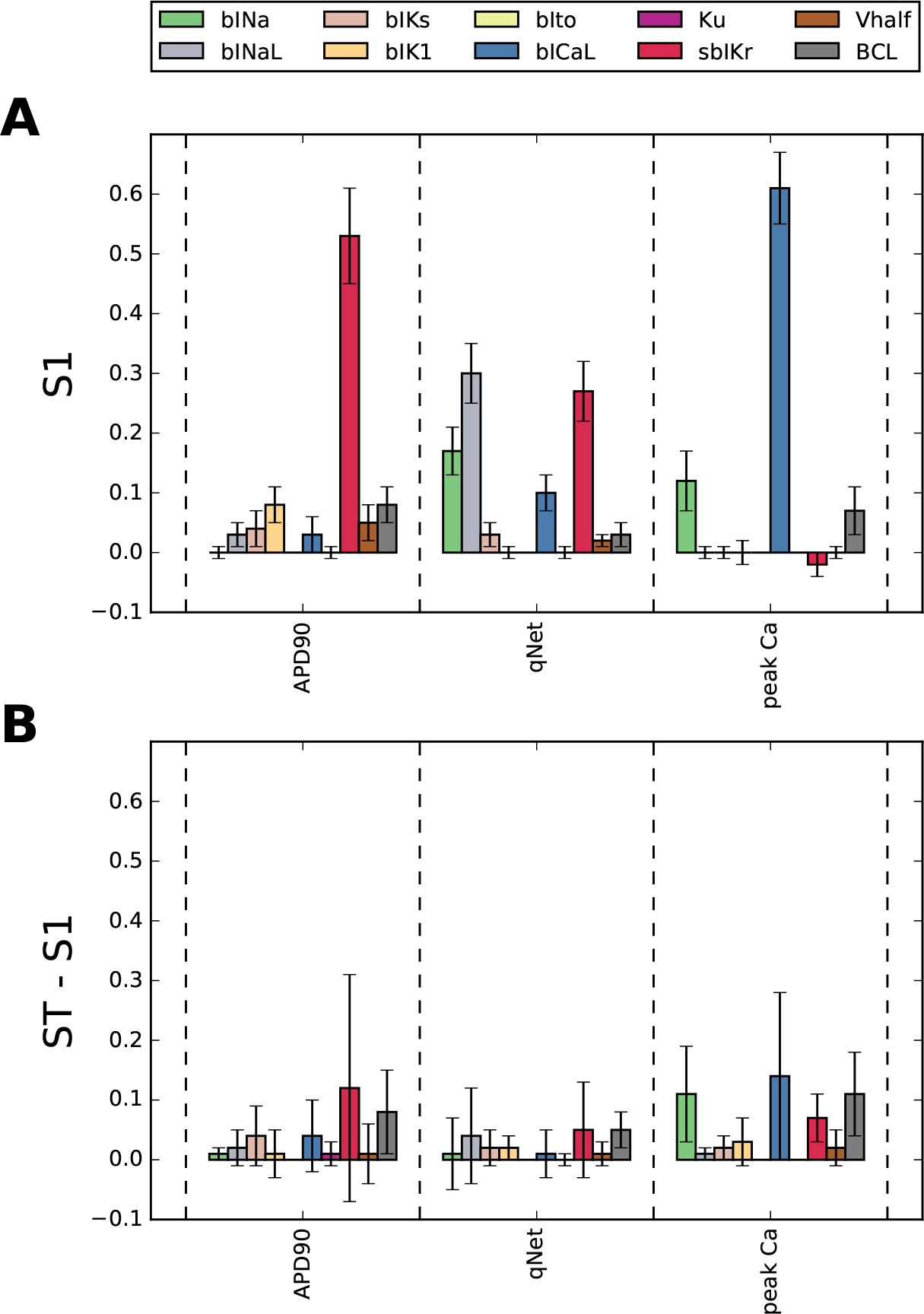
Assessment of sensitivity to blocks of different cardiac ion-channels, drug-binding parameters and *BCL* for *APD*90, *qNet*, and *peakCa* output responses in the CiPAORd endo cell model using the Sobol sensitivity indices. **A**: First-order sensitivity Sobol index, *S*1. **B**: Total sensitivity Sobol index, *ST*. The algorithm to calculate Sobol indices can produce negative values that could be eliminated by increasing sampling size.

The difference between the *ST* and *S*1 indices in Figure 3B shows the impact of higher-order indices. Small differences between *S*1 and *ST* for several derived metrics such as *APD*90, *qNet*, *peakCa* suggest minor influence of parameter interactions. The estimated sum, ∑*S*1_*i*_, of the first-order Sobol indices *S*1 for the direct features indexed by *i* (Table 2) for different model-derived metrics are listed in Table 4. ∑*S*1_*i*_ represents the contribution of all individual input parameters to the total variation of a model-derived metric without considering parameter interactions. The observed values (Table 4) indicate that *>*78% of the variance in *qNet*, *APD*90, and *peakCa* can be attributed to the individual input parameters for the endo cell model. The parameter interactions explain less than 22% of the variance of these derived metrics. Moreover, the *S*2 sensitivity index measure does not show any significant second-order interactions. This suggests that the observed small interactions effects derive from higher-order interactions terms (results are not shown). The *S*1 and *ST* sensitivity indices obtained for all the derived features (Table 3) across different cell types are reported in the Supplemental Material. The results of the sensitivity analysis using a less computationally expensive GSA method such as Morris method (elementary effects analysis) as well as multivariate linear regression methods, is also reported in the Supplemental Material for comparison. Unlike elementary effects, Sobol indices quantify the contribution of an input parameter to the metrics in isolation as well as in the presence of interaction with other parameters.

**Table 4.** Total contribution of the main effects on the variance of derived metrics estimated by the sum of the Sobol S1 index

| | $qNet$ | $APD_{90}$ | $peakCa$ |
| --- | --- | --- | --- |
| $\sum S1_i$ | 0.92 | 0.84 | 0.78 |

#### Regional sensitivity analysis

Next we wanted to determine the most influential model parameters that enhance or reduce the susceptibility to early afterdepolarizations in the CiPAORd model. To achieve this we performed Monte Carlo filtering, which is referred in literature as a method of regional sensitivity analysis. For Monte Carlo filtering, the target space was partitioned into realizations with either presence or absence of EADs in simulated action potentials under the action of a virtual drug population (*n* = 22000). Figure 4 shows the empirical CDFs for each of the 10 input parameters. The dotted and solid lines represent the estimated CDFs for the behavioral *F*_*n*_2__(*X*_*i*_|*EAD*+) and the non-behavioral *F*_*n*_1__(*X*_*i*_|*EAD*) subsets, respectively. The behavioral and non-behavioral subsets comprised 10479 and 11521 samples. If the two CDF distributions are not significantly different, than the parameter is likely unimportant, and for any value of that particular parameter in the examined range, the outputs are likely to fall into either behavioral or the non-behavioral subsets. For uniformly distributed inputs as in this study, the CDF of the non-influential parameters are close to the identity line. The bigger the distance between the empirical CDFs for the behavioral and non-behavioral subsets, the greater is the influence of the parameter to development of EADs. The figure shows that the parameter *bICaL* (Figure 4A, dashed-dotted line) has the strongest influence on susceptibility of the model to EADs. The parameters *sbIKr* (Figure 4B, solid line), *bIKs* (Figure 4C, dashed line), and *bIK*1 (Figure 4D, dashed line) have the next highest contribution to model sensitivity to EADs. The shape of the CDF curve provides additional information on the model behavior. For example, the green dashed line (Figure 4A) is steep at higher blocks of L-type calcium current. This indicates that the higher block of L-type calcium current drives the model away from EAD generation, suggesting a protective role of L-type calcium current. On the contrary, the increase in the block of *sbIKr* parameter, as expected, resulted in increased development of EADs pointing towards increased proarrhtyhmic propensity at higher blocks of hERG current.

**Figure 4.**
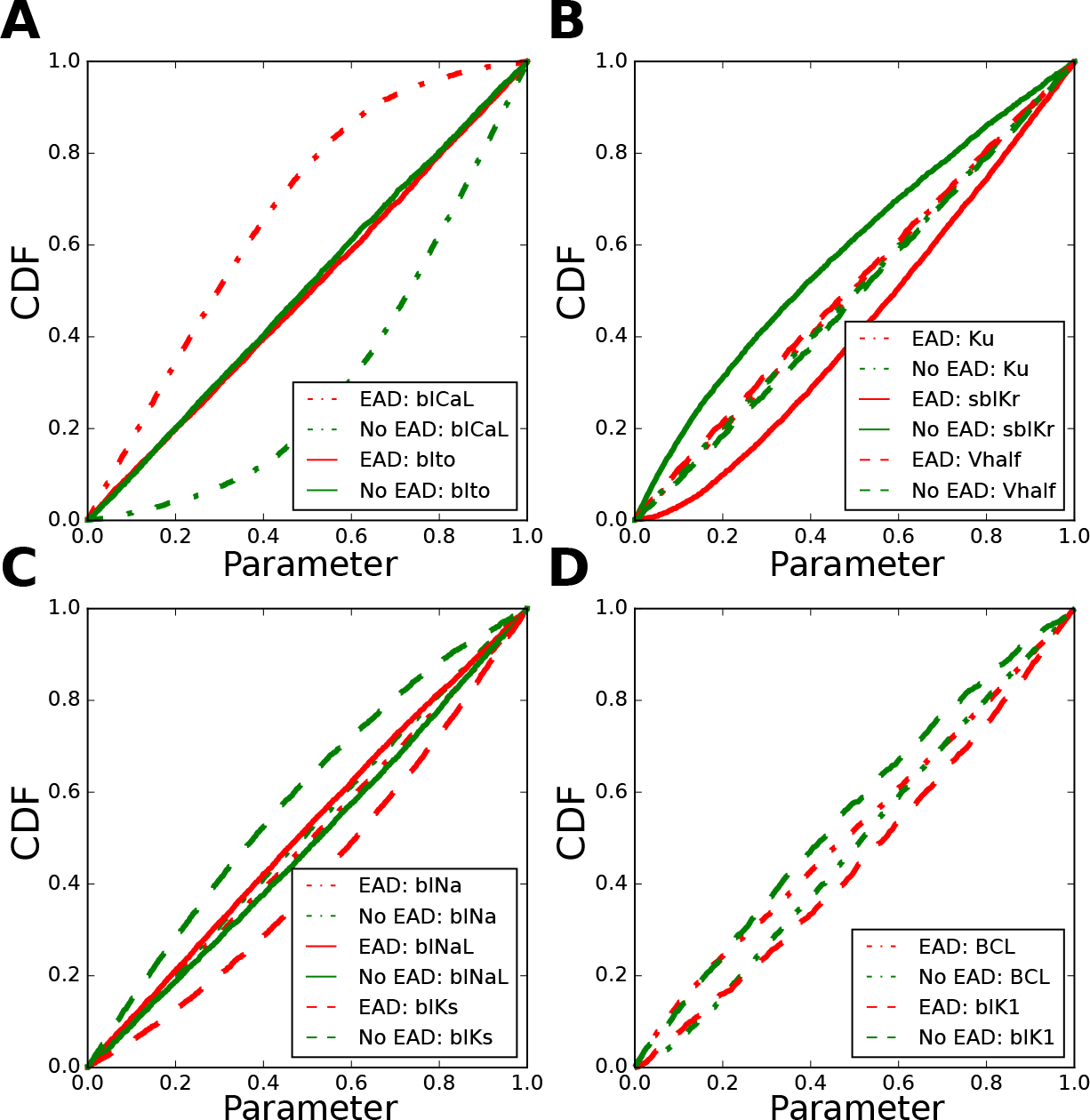
Ranking the most influential model parameters regulating EAD generation in the CiPAORd endo cell model using Monte Carlo filtering analysis. Empirical CDF for each of the 10 input parameters conditional to the presence or absence of EADs: **A**: *bIto*, *bICaL*; **B**: *Ku*, *sbIKr*, *V*_*half*_ **C**: *bINa*, *bINaL*, *bIKs*; and **D**: *bIK*1 and *BCL*.

Figure 5 provides estimates of the differences in the behavioral and non-behavioral empirical CDFs using a Kolmogorov-Smirnov two-sample test. The results show that the parameter *bICaL* regulating the block of the L-type calcium channel current had the highest influence on EAD in models with both endo and M cell types. In the M cell, the parameter *sbIKr* and *bCaL* appear to be equally important for regulation of EADs in the model. In contrast, since the additional block of the hERG channel current was required to trigger EAD in the endo cell type, the parameter *sbIKr* had moderate influence in regulation of EADs. The parameter *bIKs* had moderate influence in both endo and M cell types. The block *bIK*1 appears to have higher influence in the endo cell compared to the M cell. The parameters *Ku*, *Ito* and *bINa* were the least important. Monte Carlo filtering demonstrates that *sbIKr* and *bICaL* parameters contribute the most to generation of EADs.

**Figure 5.**
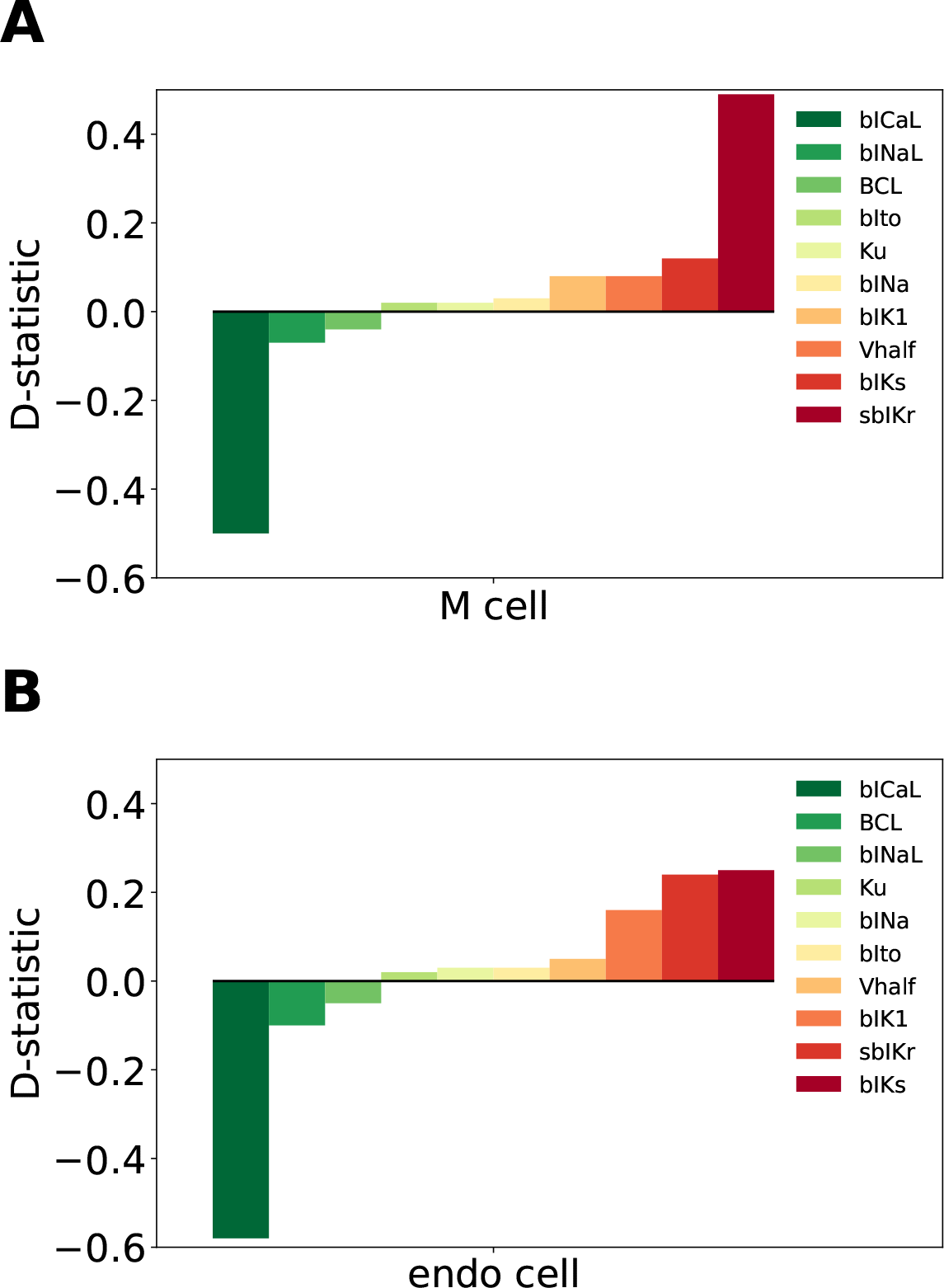
Ranking the most influential model parameters regulating EAD generation in the CiPAORd **A:** M cell and **B:** endo cell model using Monte Carlo filtering analysis. The D-statistic is obtained using Monte Carlo filtering analysis. The D-statistic obtained for the parameters with CDFs for the behavioral *F*_*n*_2__(*X*_*i*_|*EAD*+) above unity line is assigned negative signs to easily visualize if the inputs enhance or reduce the susceptibility of EADs.

#### Classification of CiPA training/validation drugs based on EADs

Here, we wanted to examine how the findings from the global sensitivity analysis on the virtual drug population would translate for a set of actual drugs with channel blocks covering only a relatively small portion of the parametric space examined previously. Specifically, we wanted to evaluate the performance of the classifiers built on the EAD feature considering only the most influential parameters as suggested by global sensitivity analysis presented in the previous sections. In addition, we also compare the performance of the classifiers based on EADs to TdP risk classifiers built on metrics such as *qNet*, which are thought to be correlated to EADs.

Figure 6 shows action potential traces obtained from simulation of 4 “CiPA compounds” using two different protocols for the endo cell CiPAORd model. In the first protocol we increased drug concentrations from 1x - 70x EFTPC to test the EAD development under different concentrations. We observed (Figure 6 A) that a drug with high torsadogenic risk like Dofetilide, resulted in EADs at relatively small concentrations (5x EFTPC). The intermediate risk drug Clarithromycin and low risk drugs Verapamil and Loratadine are not associated EADs under all the concentrations tested (Figure 6 A). We also evaluated the EAD development at a fixed drug concentration of 2x EFTPC while increasing the additional block of hERG channels from 65 to 100 % (Figure 6 B). Similar to the protocol with increased drug concentration, we observed that high risk drug Dofetilide is associated with EADs in the presence of relatively small additional perturbations of hERG current (84.5% block) compared to low and intermediate risk drugs. However, unlike the protocol with increased drug concentrations, where the compounds Clarithromycin and Loratadine are not associated with EADs at all tested concentration, presence of additional perturbations of hERG block around 94% resulted in generation of EADs for both these drugs.

**Figure 6.**
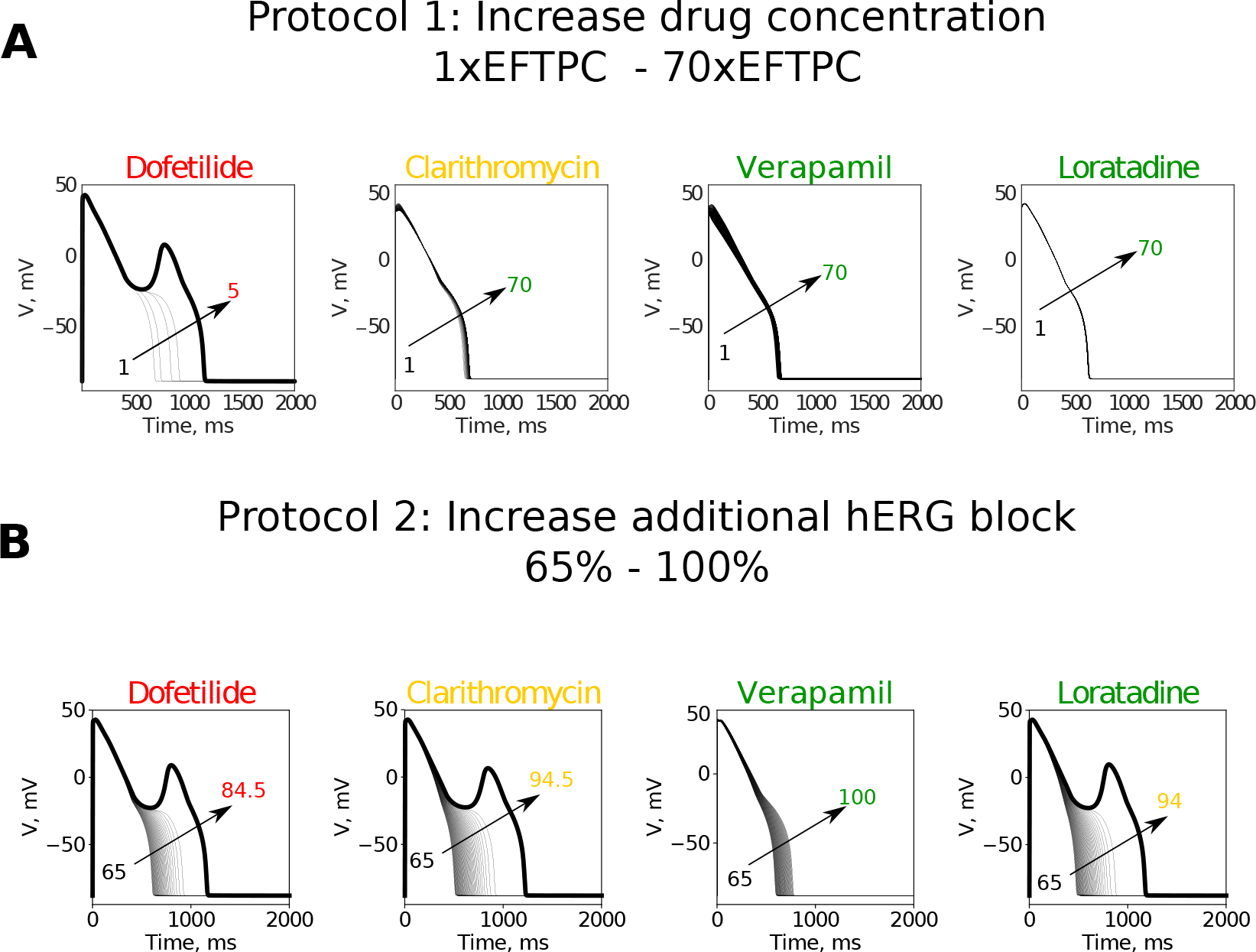
Examples of action potential transients observed at **A:** different drug concentrations at fixed hERG block of 85% and **B:** with increase of additional block of hERG channel currents at a fixed drug concentration of 2xEFPTC.

Using these two protocols we examined the classification of CiPA compounds based on the drug concentration (normalized to EFTPC) necessary to induce EADs in the CiPAORd model, defined here as *Th_EAD,conc_* and also based on the amount of additional hERG perturbation required to induce EAD in the model, denoted here as *Th_EAD,hERG_*. Since the Monte Carlo filtering results point to *sbIKr* and *bICaL* being the most critical parameters regulating EAD development, only drug-induced changes of these two parameters were taken into consideration. We also compared the obtained thresholds for EADs to the thresholds considering drug-induced changes of all the seven ion channel currents measured from the *in vitro* assays and characterized by the 9 parameters reported in Table 2. The results are summarized in the Table 5. Our EAD analysis show that the drugs in the high-risk category have consistently a threshold value of less than 7 and 90 for *Th_EAD,conc_* and *Th_EAD,hERG_*, even for the threshold obtained when considering only the drug effects on two parameters, static block of hERG channel current *sbIKr* and the block of L-type calcium channel current *bICaL*. Addition of dynamic hERG channel current parameters as well as other input parameters resulted in no significant changes in the observed thresholds for EAD generation. A high risk drug Disopyramide from the CiPA validation dataset did not induce EAD in the model under all tested conditions. The intermediate risk drugs Cisapride and Ondansetron resulted in EADs in the model at a threshold of less than 7 similar to the drugs in high risk category under all tested conditions. Similarly, Ranolazine and Metoprolol drugs that are defined as low risk under the CiPA initiative had a threshold value of less than 7 and 90 for *Th_EAD,conc_* and *Th_EAD,hERG_*, respectively, for all the conditions tested. The low-risk drugs Diltiazem, Mexiletine, Verapamil, Loratadine, Nifedipine, and Nitrendipine did not produce EADs in the model under all the tested conditions for the protocol with increase in drug concentrations. Similar results were observed for the additional hERG perturbation protocol except for the drugs Loratadine and Mexiletine. Chlorpromazine resulted in EADs at relatively high threshold compared to the high risk drugs with a threshold of *>* 14 and *>* 91 for *Th_EAD,conc_* and *Th_EAD,hERG_*, respectively, under all conditions. Low risk drug Tamoxifen consistently resulted in EADs in the model at thresholds values similar to intermediate risk drugs. Terfenadine, Pimozide, and Clozapine were the only few drugs with significant changes in the observed threshold and switching to a different risk category when the drug-induced changes of parameters other than *sbIKr* and *bICaL* were ignored for the protocol with increased drug concentration. Similarly, few drugs like Droperidol, Pimozide, Mexiletine and Terfenadine switched risk category when the drug-induced changes of parameters other than *sbIKr* and *bICaL* were not considered for the protocol with additional hERG perturbation.

**Table 5.** Thresholds of EAD (xEFTPC and hERG channel pertrubation) and *qNet* for CiPA training (12 drugs) and validation (16 drugs) datasets

| Drug | $Th_{EAD,conc}$ | | | | $Th_{EAD,hERG}$ | | $qNet$ | TdP risk |
| --- | --- | --- | --- | --- | --- | --- | --- | --- |
|  | endo cell |  | M cell |  | endo cell |  | endo cell |  |
|  | C1 | C2 | C1 | C2 | C1 | C2 | C1 |  |
| Quinidine | 1 | 1 | 1 | 1 | 65.0 | 74.0 | 12.56 | High |
| Ibutilide | 1 | 1 | 1 | 1 | 65.0 | 65.0 | 6.71 | High |
| Azimilide | 2 | 4 | 3 | 4 | 83.0 | 87.5 | 49.90 | High |
| Bepridil | 3 | 2 | 3 | 2 | 84.5 | 79.0 | 45.64 | High |
| Dofetilide | 3 | 3 | 6 | 4 | 84.5 | 86.0 | 57.12 | High |
| Vandetanib | 3 | >70 | 3 | 4 | 90.0 | 90.0 | 49.40 | High |
| Sotalol | 4 | 5 | 5 | 6 | 87.5 | 88.5 | 57.32 | High |
| Ranolazine | 4 | 3 | 5 | 3 | 89.0 | 83.0 | 73.92 | Low |
| Cisapride | 5 | 2 | 7 | 3 | 87.5 | 83.0 | 61.43 | Medium |
| Metoprolol | 6 | 5 | 6 | 6 | 90 | 89.0 | 58.02 | Low |
| Ondansetron | 6 | 7 | 7 | 7 | 89.5 | 90.0 | 63.13 | Medium |
| Astemizole | 9 | 9 | 22 | 12 | 91.0 | 91.5 | 65.36 | Medium |
| Droperidol | 10 | 7 | 25 | 10 | 90.5 | 89.5 | 62.48 | Medium |
| Chlorpromazine | 14 | 35 | 21 | 27 | 91.5 | 92.5 | 66.42 | Medium |
| Terfenadine | 17 | 4 | 15 | 4 | 90.5 | 87.5 | 60.49 | Medium |
| Tamoxifen | 17 | 31 | 20 | 27 | 93.5 | 93.5 | 69.73 | Low |
| Nitrendipine | >70 | >70 | >70 | >70 | 100 | 100 | 77.53 | Low |
| Pimozide | >70 | 4 | 20 | 3 | 92.5 | 88.0 | 62.6 | Medium |
| Clozapine | >70 | >70 | 23 | >70 | 93.0 | 93.0 | 68.17 | Medium |
| Risperidone | >70 | >70 | >70 | >70 | 94.0 | 93.5 | 70.14 | Medium |
| Clarithromycin | >70 | >70 | >70 | >70 | 94.5 | 94.5 | 69.34 | Medium |
| Diltiazem | >70 | >70 | >70 | >70 | 100 | 100.0 | 88.59 | Low |
| Mexiletine | >70 | >70 | >70 | >70 | 100 | 96.5 | 88.97 | Low |
| Verapamil | >70 | >70 | >70 | >70 | 100 | 100.0 | 73.99 | Low |
| Disopyramide | >70 | >70 | >70 | >70 | 95.5 | 95.5 | 72.03 | High |
| Loratadine | >70 | >70 | >70 | >70 | 94.0 | 94.0 | 70.30 | Low |
| Domperidone | >70 | >70 | >70 | >70 | 100 | 100.0 | 58.45 | Medium |
| Nifedipine | >70 | >70 | >70 | >70 | 100 | 100.0 | 84.97 | Low |

**TdP risk classification summary**
| Category | No. correctly classified drugs |  |  |  | No. correctly classified |  | No. correctly classified | Total number of Drugs |
| --- | --- | --- | --- | --- | --- | --- | --- | --- |
|  | C1 | C2 | C1 | C2 | C1 | C2 | C1 |  |
| High | 7 (4, 3) | 6 (4, 2) | 7 (4, 3) | 7 (4, 3) | 7 (4, 3) | 7 (4, 3) | 7 (4, 3) | 8 (4, 4) |
| Intermediate | 4 (2, 2) | 2 (1, 1) | 6 (2, 4) | 3 (1, 2) | 8 (2, 6) | 5 (1, 4) | 11 (4, 7) | 11 (4, 7) |
| Low | 6 (3, 3) | 6 (3, 3) | 6 (3, 3) | 6 (3, 3) | 5 (3, 2) | 4 (2, 2) | 6 (4, 2) | 9 (4, 5) |
| Total | 17 (9, 8) | 14 (8, 6) | 19 (9, 10) | 16 (8, 8) | 20 (9, 11) | 16 (7, 9) | 24 (12, 12) | 28 (12, 16) |

<sup>C1</sup> Drug-induced modulation of 9 parameters ( $sbIKr$ , $Ku$ , $V_{half}$ , $bINa$ , $bINaL$ , $bICaL$ , $bIKs$ , $bIK1$ and $bIto$ ) is considered.
<sup>C2</sup> Only drug-induced changes in $sbIKr$ and $bICaL$ is considered ( $V_{half} = -100$ , $Ku = 0.05$ ).
Note Numbers in parentheses are number of drug from training and validation set.
Note BCL fixed to 2000 for endo cell type and 700 for M cell type model under all tested conditions.

Although EADs are thought to be cellular precursors of TdP, the classifier based on EADs alone ranks correctly only 17-20 drugs, thus performing worse than *qNet*. In the table (Table 5) we also report the estimated *qNet* values for the 28 drugs at 4x EFTPC drug concentrations. We observe the drugs like Ranolazine, Cisapride, Ondansetron, and Domperidone, which are not correctly classified by either of the EAD based classifiers, are correctly classified by *qNet*.

## 4 DISCUSSION

Uncertainties in *in vitro* measurements of drug-induced effects on ionic currents present an important concern in evaluating the torsadogenic risk of compounds by interrogating *in silico* biophysical models. Discrepancies in estimates for model parameters based on available *in vitro* assay data have been recently highlighted in uncertainty quantification studies (Johnstone et al., 2016; Chang et al., 2017). High uncertainty in model parameters leads to low confidence in model predicted risk, and thus, not surprisingly, risk stratification of the CiPA training drugs proved to be unreliable especially at high drug concentrations (Chang et al., 2017), where model parameter estimates are inherently less accurate. However, it is important to emphasize that the relative contributions of drug-induced modulation of ion-channels on output features differ significantly. Uncertainties in model input parameters that are highly influential (e.g., as revealed by sensitivity analysis) result, therefore, in lower confidence in the predicted risk, while errors in estimating less influential model parameters are better tolerated by risk measures (Loucks et al., 2017; Mirams et al., 2016). In this paper, we present a study that applies GSA within the context of *in silico* prediction of pharmacological toxicity. The target of GSA was the latest version of the *in silico* model of an isolated cardiac cell (Dutta et al., 2017), CiPAORd, which was developed under the CiPA initiative and incorporates dynamic hERG-drug interactions (Li et al., 2017). Our analysis explored the effects of a large population of virtual drugs on the seven major cardiac ion-channel currents thought to be important in regulation of TdP. GSA provided a systematic understanding of the model input-output relationships and allowed for the identification of the most influential parameters that regulate model-derived features used for proarrhythmic risk classification. The knowledge gained from GSA could help further improve the model structure and increase reliability of model-predicted risk.

### Sensitivity analysis used for cardiac models

Different methods and tools, each with their own advantages and disadvantages, allow for the analysis of the sensitivity of complex systems to the input parameters (e.g., refer to (Saltelli et al., 2008; Iooss and Lemaître, 2014; Pianosi et al., 2016) for thorough reviews on the subject). Simple sensitivity analyses performed by varying one parameter at a time have been carried out to asses the impact of changes in ionic currents on cardiac cellular electrophysiology (Romero et al., 2009; Chang et al., 2014). This type of sensitivity analysis, although computationally inexpensive, only quantifies the impact on model outputs of changes in a single input parameter relative to the point estimates chosen for the rest of the parameters that are held constant. On the contrary, GSA quantifies the effects of global variations over the entire input parameter space. Multivariate linear regression models that rely on AAT sampling approaches have been used in the past on the cardiac cellular models (Sobie, 2009) to identify how changes in model parameters affect different outputs of the model, to address different physiological questions, to improve model structure, and to suggest novel experiments (Cummins et al., 2014; Sarkar and Sobie, 2010; Britton et al., 2013; Sadrieh et al., 2013; Devenyi et al., 2017; Devenyi and Sobie, 2016; Lee et al., 2013). Recently, application of multivariate logistic regression has been reported to relate perturbations in model parameters to the presence/absence of EADs (Morotti and Grandi, 2016). The multivariate linear regression is suitable and accurate for models with almost linear input-output relationship. Similarly, the logistic regression applied to determine EAD sensitivity is accurate if a surface separating EAD and non-EAD regions is close to a hyperplane.

### Critical inputs regulating *APD*90, *qNet*, *peakCa*

In this study we applied a more general form of GSA that is suitable even in presence of non-linear input-output relationships (Saltelli et al., 2008). In particular we used the Sobol variance-based sensitivity method (Saltelli et al., 2008; Sobol’, 2001) to rank cardiac ion-channel currents based on their relative contributions to variability in the model-derived features. We also performed sensitivity analysis to determine the cardiac ion-channels that regulate EAD generation in the CiPAORd models using Monte Carlo filtering methods Hornberger and Spear (1981); Saltelli et al. (2008). Our systematic sensitivity analysis identified critical input parameters for the variability of the different model-derived features used for TdP risk assessment (see Figure 3 and data in the Supplemental Material). More specifically, we observed that the recently proposed *qNet* metric is most sensitive to modulations in sodium currents and to the *sbIKr* parameter. *sbIKr*, *bIK*1 and *bICaL* were found to be the most influential parameters regulating *APD*90 (Figure 3). In the past, *APD*90 has also been shown, by varying one parameter at a time in the original ORd model (O’Hara et al., 2011), to be most sensitive to block of hERG current. Furthermore, the QT interval measured in 3D human-heart simulations (Sahli Costabal et al., 2019) with original ORd model (O’Hara et al., 2011) at the cellular level exhibits similar sensitivity profile as *APD*90. This is in agreement with previous observations of high correlation between *APD*90 and QT interval in the cardiac model simulations (Beattie et al., 2013). In our study, features derived from the calcium transient such as *peakCa* were found, as expected, to be most sensitive to the *bICaL* parameter.

In spite of the observed differences in the sensitivity profiles, different derived metrics have been reported to perform well on certain *in vitro* datasets. *APD*90 (Mirams et al., 2011), a metric based on *APD*50 and *diastolicCa* (Lancaster and Sobie, 2016), and a metric based on EADs (Christophe, 2013) have been shown to provide good risk discrimination of drugs considering *in vitro* measurements reported in (Mirams et al., 2011). We have shown previously that different derived features extracted from the original ORd model (O’Hara et al., 2011) show similar performance in TdP risk discrimination (Parikh et al., 2017) when tested on a combination of several datasets. The similarity in performance of different metrics might be due to the presence of estimates of drug effects on only three channel currents (i.e., fast sodium current, L-type calcium channel current, and hERG current) in the majority of the datasets, the small size of the datasets, and the differences in structure of the myocyte models used for obtaining the derived feature. Different derived metrics, such as *APD*50, *APD*90, *peakCa*, and *CaD*90 have been shown to provide the best classification when varying the computational model of interest (Mirams et al., 2011).

Several cardiac ion-channel/parameters that are thought to be important for improved drug-induced TdP risk assessments and measured experimentally via *in vitro* ion-channel screening (Crumb et al., 2016) showed really minor influence in regulation of the model-derived features. For example, the block of transient outward current and the dynamic hERG block parameters showed relatively minor influence on majority of the tested metrics. Specifically *qNet* metrics appeared to be insensitive to the *bIK*1, *bto* and *Ku* parameters (Figure 3 and the Supplemental Material).

GSA results are highly dependent on explored parametric space. Here, we evaluate the sensitivity over the 10-D input space comprising parameters of seven major cardiac channel currents that are thought to play an important role in determining the risk of TdP. However, the actual drugs might lie within a very small subspace of the explored hyperspace. Visualization of the blocks of different ion channel currents for the 28 CiPA drugs (Figure 7) reveals that majority of the drugs do not result in block of *IK*1, *INa*, and *IKs* currents. Figure 7 demonstrates that accurate classification of the Ranolazine drug to low risk category requires a feature that is at least moderately sensitive to variations in block of late sodium current, since the drug appears to be a pure hERG and sodium channel blocker. Our GSA results (Figure 3) point to *qNet* as a candidate, as it is the only feature among the tested derived features that is highly sensitive to block of late sodium current and block of hERG. *qNet* has been observed to outperform other standard derived features on the 28 CiPA drugs (Dutta et al., 2017; Li et al., 2018, 2017).

**Figure 7.**
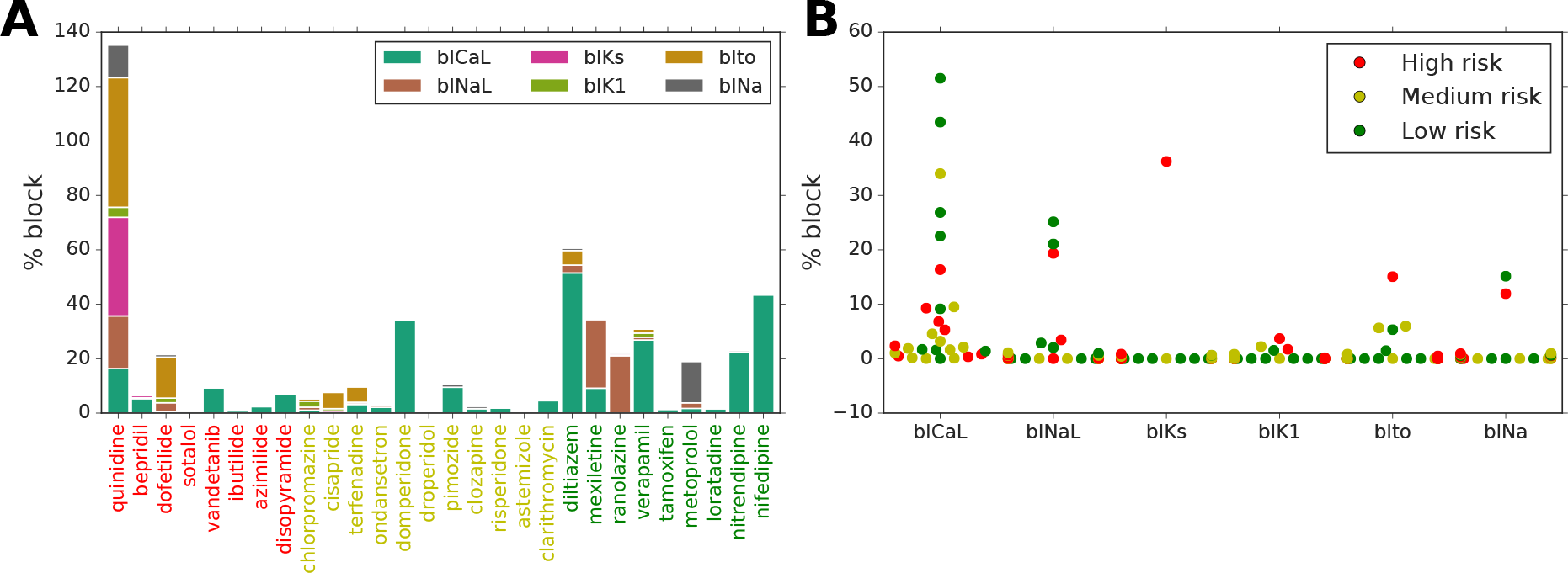
Drug-induced block of non-hERG ion channels for 28 CiPA compounds at their EFTPC based on the measurements from the *in vitro* assay Crumb et al. (2016); Li et al. (2018). **A**: Stacked bar chart of six ion channel current block values for each of the 28 drugs.**B**: A swarm plot of block values of six ion channel currents categorized into high, medium, and low risk groups.

### EAD sensitivity analysis

The EAD sensitivity analysis (Figure 4 and 5) indicates that the generation of EADs, which are thought to be cellular precursors of TdP genesis (Yan et al., 2001), are most sensitive to variations in block of *ICaL* and to the static component of the hERG channel current in the CiPAORd model. Block of hERG channels is well established to be critical for generation of EADs and eventually Torsades de Pointes (Redfern et al., 2003) and has been shown to be the primary current responsible for generation of EADs in the simulations using original ORd model (Christophe, 2013). The role of L-type calcium channel currents in regulation of EADs have been also highlighted across different studies (January and Riddle, 1989; Zeng and Rudy, 1995; Weiss et al., 2010). Our results show that variations in blocks of the *IKs* and *IK*1 currents have a moderate influence on the genesis of EADs in the CiPAORd model. Drug effects on the *Ito*, *INa* have the least influence on EAD generation (Figure 5 and 4). The recently introduced dynamic-hERG block parameters *V*_*half*_ and *Ku* (Li et al., 2017), which are measured using challenging experimental protocols (Milnes et al., 2010; Veroli et al., 2014), show minor influence on EAD genesis (Table 5) when compared to other tested inputs. These parameters also exhibit relatively small contribution to the variance of all the tested derived metrics compared to other influential input parameters (Figure 3, and data in the Supplemental Material). However, it should be noted that in cases where the majority of the primary regulating parameters are similar between drugs, accounting for changes in the modestly influential parameters can allow for improved predictions. On classifying CiPA drugs based on EADs, we observed that prediction improves by correctly classifying 3 more drugs when accounting for drug-induced effects of other parameters in addition to the *sbIKr* and *bICaL* parameters (Table 5). However, our results also point towards the important consideration that errors in measuring the most influential parameters regulating a particular metric have a bigger impact on the predicted classification compared to neglecting some of the less influential parameters. GSA allows us to determine and rank most of the critical model components.

### Mechanistic insight from model-derived metrics

Simple statistical classifiers based on direct feature from our group and others have been shown previously to provide equivalent performance as biophysically detailed models for TdP (Mistry, 2018; Kramer et al., 2013; Mistry et al., 2015; Parikh et al., 2017). Our sensitivity analysis results also highlight strong linearity between the inputs and different model-derived metrics (such as *qNet*, *APD*90, etc.) that are proposed for TdP risk stratification (Table 4). The metric linearity suggests that the model-derived metrics can be well captured as a linear combination of the set of direct features and provides a plausible explanation for equivalent performance of the simple statistical methods. Almost linear input-output relationship in different cardiac models has also been observed in several previous studies (Sobie, 2009; Sarkar and Sobie, 2010). However, one of the most appealing feature for the biophysical models is that of interpretability, i.e. the model-derived features attempt to capture the aspects of the underlying physiological phenomena such as APD prolongation or increase in calcium levels to provide a mechanism-based classifier. Being biophysically motivated, classifiers built on model-derived features are thought to allow generalizable assessments also in cases where the training datasets are small and hence the effects on targets of interest might need to be extrapolated. A promising metric *qNet*, proposed by the modeling team at FDA (Dutta et al., 2017), has recently been shown to provide excellent classification of drugs in the CiPA training and validation data, a result thought to be linked to EAD generation (Dutta et al., 2017; Li et al., 2018). However, our GSA results show that none of the tested derived features demonstrating identical sensitivity profile to EAD (Figure 3, 5 and the Supplemental material). The *qNet* metric was observed to be sensitive to variations in block of sodium currents and block of *sbIKr* for the endo cell model. In contrast, the *bICaL* and *sbIKr* parameters are found to be the most influential for EADs. Moreover, we observed that the categorization of CiPA drugs based on analysis of EADs was not as predictive as model-derived metric such as *qNet* (Table 5). We found that drugs like Ranolazine, Cisapride, Ondansetron, and Domperidone, which were not correctly classified by either of the EAD based classifiers, were correctly classified by *qNet*. Hence, the previously reported correlation between *qNet* and EAD generation (Dutta et al., 2017) seems not to be well justified, and our results highlight the need of further research to better understand the mechanistic underpinning of the success of this promising metric. One possible speculation for improved predictive power of *qNet* for drugs like Ranolazine might be the reduced transmural dispersion of repolarization (Shimizu and Antzelevitch, 1998), which is affected by the block of the late sodium current. There can be several other possible explanations for poor performance of EAD metric compared to *qNet*, such as inaccurate reproduction of EADs in the current model, small size of the tested datasets, biases in the target, and the need to test EADs on coupled cells/tissue models.

### Summary

The proarrhythmic risk assessment based on simulated drug responses in isolated cell model (Mirams et al., 2011; Christophe, 2013, 2015; Lancaster and Sobie, 2016; Li et al., 2017; Dutta et al., 2017; Passini et al., 2017; Li et al., 2018; Parikh et al., 2017), tissue models (Kubo et al., 2017; Trenor et al., 2013) or organ level computational models (Okada et al., 2015; Costabal et al., 2018; Sahli Costabal et al., 2019) provide important physiological and mechanistic insights. Moreover, *in silico* models serve as an excellent tool for evaluation of drug safety in diseased conditions (Trenor et al., 2013; Kubo et al., 2017). However, the uncertainties in pharmacological data used for model-driven predictions and in the intrinsic structures of biophysical models used for cardiotoxic risk predictions present fundamental challenges. In this study, we showed potential application of sensitivity analysis for improved model-based proarrhythmic risk predictions. The critical model inputs regulating the model-derived metrics such as *APD*90 and *qNet* proposed for evaluation of proarrhythmic risk were identified. The analysis highlighted the need for better mechanistic understanding of promising metrics such as *qNet* and provided possible explanation for equivalent performance of the simple statistical based-classifiers and complex model-driven risk predictions. In conclusion, the sensitivity analysis method in addition with uncertainty quantification approaches can form an important component of the model-based cardiotoxic risk prediction pipeline. An improved pipeline would ultimately allow for refinement of existing biophysical models to achieve increased confidence in the model-driven proarrhtymic risk predictions.

## AUTHOR CONTRIBUTIONS

JP designed the study, performed simulations, analyzed results and wrote the manuscript. PD designed the study, analyzed the results and wrote the manuscript. JK wrote the manuscript and supervised the project. VG designed the study, analyzed the results, wrote the manuscript and supervised the project. All authors agree to be accountable for the content of the work.

## DATA AVAILABILITY STATEMENT

The datasets analyzed for this study can be found in the supplemental material.

## Footnotes

1 Here and further in the paper, we discuss linear regressions with input features typically used in the sensitivity analysis of cell models, i.e., regressions with only linear combinations of features constructed from the input parameters.

