## Supplemental Material for "Intrinsic structure of model-derived metrics for *in silico* proarrhytmic risk assessment identified by global sensitivity analysis"

### Supplementary Material

#### 1 SUPPLEMENTARY DATA

##### CiPA training and validations compounds dataset

$IC_{50}$ values and Hill coefficients describing the effects of the CiPA training drugs on non-hERG channels were extracted from Li et al. (2017). Similarly, median values from the manual patch clamp experiments on the CiPA validation drugs were taken from Li et al. (2018). The values are reported in Table S1.

**Table S1.** $IC_{50}$ values (nM) and Hill coefficients for non-hERG channels

| | ICaL_IC50 | ICaL_h | IK1_IC50 | IK1_h | IKs_IC50 | IKs_h | INa_IC50 | INa_h | INaL_IC50 | INaL_h | Ito_IC50 | Ito_h | hERG_IC50 | hERG_h |
|---|---|---|---|---|---|---|---|---|---|---|---|---|---|---|
| astemizole | 553.0 | 1.2000 | - | - | - | - | 5.410000e+03 | 0.7600 | 10300.0 | 2.3000 | - | - | - | - |
| azimilide | 13200.0 | 0.7100 | - | - | - | - | 3.630000e+05 | 0.7200 | 2940000.0 | 0.4700 | - | - | - | - |
| bepridil | 2808.0 | 0.6486 | - | - | 28630.0 | 0.7061 | 2.929000e+03 | 1.1640 | 1814.0 | 1.4160 | 8.594000e+03 | 3.5410 | 51.340 | 0.9293 |
| chlorpromazine | 8192.0 | 0.8441 | 9270.0 | 0.6878 | - | - | 4.536000e+03 | 1.9950 | 4560.0 | 0.9379 | 1.762000e+07 | 0.3654 | 975.200 | 0.8281 |
| cisapride | 9267000.0 | 0.4261 | 29480.0 | 0.5133 | 81170000.0 | 0.2921 | - | - | - | - | 2.191000e+05 | 0.2430 | 11.270 | 0.6210 |
| clarithromycin | 38100.0 | 0.8800 | - | - | - | - | 1.090000e+06 | 0.8900 | 1810000.0 | 3.0000 | - | - | - | - |
| clozapine | 5490.0 | 0.9400 | - | - | - | - | 2.570000e+05 | 0.5900 | 73600.0 | 2.0000 | - | - | - | - |
| diltiazem | 112.1 | 0.7142 | - | - | - | - | 1.109000e+05 | 0.7022 | 21870.0 | 0.6779 | 2.822000e+09 | 0.1696 | 13140.000 | 0.9119 |
| disopyramide | 32900.0 | 0.6900 | - | - | - | - | 1.920000e+05 | 1.3000 | 377000.0 | 2.1000 | - | - | - | - |
| dofetilide | 260.3 | 1.1630 | 394.4 | 0.7650 | - | - | 3.805000e+02 | 0.8920 | 753100.0 | 0.2597 | 1.882000e+01 | 0.7712 | 6.125 | 1.0790 |
| domperidone | 73.6 | 0.4900 | - | - | - | - | 4.190000e+04 | 1.5000 | 225000.0 | 2.1000 | - | - | - | - |
| droperidol | 3230.0 | 1.2000 | - | - | - | - | 3.660000e+04 | 2.5000 | 33800.0 | 2.8000 | - | - | - | - |
| ibutilide | 37000.0 | 0.8600 | - | - | - | - | 2.410000e+04 | 2.3000 | 287000.0 | 2.4000 | - | - | - | - |
| loratadine | 703.0 | 0.5600 | - | - | - | - | 1.130000e+05 | 1.4000 | 192000.0 | 2.1000 | - | - | - | - |
| metoprolol | 3280000.0 | 0.5400 | - | - | - | - | 3.030000e+04 | 0.6100 | 630000.0 | 0.6600 | - | - | - | - |
| mexiletine | 38240.0 | 1.0310 | - | - | - | - | - | - | 8957.0 | 1.4090 | - | - | 29070.000 | 0.8928 |
| nifedipine | 11.4 | 0.6700 | - | - | - | - | 2.760000e+04 | 1.1000 | 45600.0 | 4.5000 | - | - | - | - |
| nitrendipine | 35.7 | 0.5000 | - | - | - | - | 2.240000e+04 | 0.5800 | 70700.0 | 3.2000 | - | - | - | - |
| ondansetron | 22550.0 | 0.7526 | - | - | 569800.0 | 0.6535 | 5.767000e+04 | 1.0200 | 19180.0 | 1.0350 | 1.023000e+06 | 0.9891 | 1325.000 | 0.9210 |
| pimozide | 64.5 | 0.4500 | - | - | - | - | 1.020000e+04 | 0.4600 | 1910.0 | 1.9100 | - | - | - | - |
| quinidine | 51590.0 | 0.5892 | 39590000.0 | 0.3468 | 4899.0 | 1.3630 | 1.233000e+04 | 1.4940 | 9417.0 | 1.3370 | 3.487000e+03 | 1.2820 | 986.300 | 0.8404 |
| ranolazine | - | - | - | - | 36160000.0 | 0.5191 | 6.877000e+04 | 1.4250 | 7884.0 | 0.9450 | - | - | 8208.000 | 0.8576 |
| risperidone | 1470.0 | 0.5900 | - | - | - | - | 5.340000e+05 | 0.7100 | 11400000.0 | 5.8000 | - | - | - | - |
| sotalol | 7062000.0 | 0.8651 | 3050000.0 | 1.2040 | 4222000.0 | 1.1670 | 1.144000e+09 | 0.5089 | - | - | 4.314000e+07 | 0.6632 | 107100.000 | 0.7850 |
| tamoxifen | 5720.0 | 0.7600 | - | - | - | - | 8.400000e+04 | 0.8200 | 3640000.0 | 4.0000 | - | - | - | - |
| terfenadine | 700.4 | 0.6601 | - | - | 399800.0 | 0.5430 | 4.803000e+03 | 1.0150 | 20060.0 | 0.6011 | 2.400000e+05 | 0.2559 | 20.400 | 0.6118 |
| vandetanib | 6060.0 | 0.7200 | - | - | - | - | 8.090000e+04 | 1.9000 | 3790000.0 | 0.8200 | - | - | - | - |
| verapamil | 201.8 | 1.0970 | 348800000.0 | 0.2728 | - | - | - | - | 7028.0 | 1.0310 | 1.343000e+04 | 0.8222 | 295.600 | 0.9378 |

hERG dynamic data was extracted from Li et al. (2017) and Li et al. (2018) for the training and validation compounds, respectively, as listed in Table S2. In the simulations, the reported median values of parameters were used.

**Table S2.** Drug-hERG binding dynamic parameters.

| | Kmax | Ku | n | halfmax | Vhalf |
|---|---|---|---|---|---|
| astemizole | 2.420 | 0.000033 | 1.4490 | 4.883000e+00 | -6.110 |
| azimilide | 654000.000 | 0.008250 | 0.6028 | 1.413000e+07 | -8.821 |
| bepridil | 5594000.000 | 0.000172 | 0.9374 | 1.472000e+08 | -61.340 |
| chlorpromazine | 157900.000 | 0.046710 | 0.8871 | 4.351000e+07 | -14.450 |
| cisapride | 10.220 | 0.000416 | 0.9615 | 4.232000e+01 | -167.400 |
| clarithromycin | 92.890 | 0.012160 | 0.7867 | 2.229000e+05 | -102.800 |
| clozapine | 7.486 | 0.029890 | 1.3670 | 9.048000e+04 | -8.810 |
| diltiazem | 182500.000 | 0.282000 | 0.9382 | 6.677000e+08 | -90.650 |
| disopyramide | 3.685 | 0.121600 | 0.7894 | 4.473000e+04 | -78.110 |
| dofetilide | 35.100 | 0.000018 | 1.0800 | 2.166000e+02 | -1.000 |
| domperidone | 3.339 | 0.000356 | 0.7026 | 1.609000e+01 | -65.650 |
| droperidol | 14.210 | 0.001256 | 0.5780 | 1.165000e+02 | -78.680 |
| ibutilide | 14.570 | 0.000061 | 0.9231 | 3.863000e+01 | -9.771 |
| loratadine | 376500.000 | 0.009642 | 0.8368 | 4.770000e+08 | -1.000 |
| metoprolol | 31790.000 | 0.850800 | 0.8110 | 4.223000e+07 | -89.350 |
| mexiletine | 15.000 | 0.071140 | 1.1390 | 7.230000e+05 | -87.510 |
| nifedipine | 4.748 | 0.991600 | 1.2350 | 9.752000e+08 | -87.370 |
| nitrendipine | 1.713 | 0.981900 | 1.9230 | 5.669000e+08 | -61.580 |
| ondansetron | 172000.000 | 0.023240 | 0.8910 | 5.224000e+07 | -82.200 |
| pimozide | 10.070 | 0.000046 | 0.8714 | 5.601000e+00 | -158.500 |
| quinidine | 275.700 | 0.004103 | 0.8488 | 5.383000e+04 | -61.350 |
| ranolazine | 52.840 | 0.020350 | 0.9532 | 1.430000e+05 | -94.990 |
| risperidone | 3.930 | 0.001151 | 1.1220 | 7.528000e+02 | -80.430 |
| sotalol | 96190.000 | 0.022250 | 0.7513 | 3.856000e+08 | -51.500 |
| tamoxifen | 3.900 | 0.011750 | 2.0000 | 4.067000e+05 | -2.036 |
| terfenadine | 102200.000 | 0.000078 | 0.6502 | 4.095000e+05 | -81.630 |
| vandetanib | 36.280 | 0.019740 | 0.7126 | 2.223000e+03 | -48.550 |
| verapamil | 1694000.000 | 0.000816 | 1.0430 | 3.356000e+08 | -97.080 |

The known TdP categories and maximum effective free therapeutic concentrations for the validation drugs (Li et al., 2017, 2018) are listed in Table S3.

**Table S3.** Drug Therapeutic concentrations and risk

| Drug | EFTPC (nM) | Risk |
|---|---|---|
| astemizole | 0.260 | Medium |
| azimilide | 70.000 | High |
| bepridil | 33.000 | High |
| chlorpromazine | 38.000 | Medium |
| cisapride | 2.600 | Medium |
| clarithromycin | 1206.000 | Medium |
| clozapine | 71.000 | Medium |
| diltiazem | 122.000 | Low |
| disopyramide | 742.000 | High |
| dofetilide | 2.000 | High |
| domperidone | 19.000 | Medium |
| droperidol | 6.330 | Medium |
| ibutilide | 140.000 | High |
| loratadine | 0.450 | Low |
| metoprolol | 1800.000 | Low |
| mexiletine | 4129.000 | Low |
| nifedipine | 7.700 | Low |
| nitrendipine | 3.020 | Low |
| ondansetron | 139.000 | Medium |
| pimozide | 0.431 | Medium |
| quinidine | 3237.000 | High |
| ranolazine | 1948.200 | Low |
| risperidone | 1.810 | Medium |
| sotalol | 14690.000 | High |
| tamoxifen | 21.000 | Low |
| terfenadine | 4.000 | Medium |
| vandetanib | 255.400 | High |
| verapamil | 81.000 | Low |

#### Global sensitivity analysis results

##### Visulation of changes in the model-derived metrics against individual input parameters

The figures (Figure S1, S2) show values of the derived features extracted from the simulation of virtual drugs generated for the Sobol sensitivity analysis against individual input parameters.

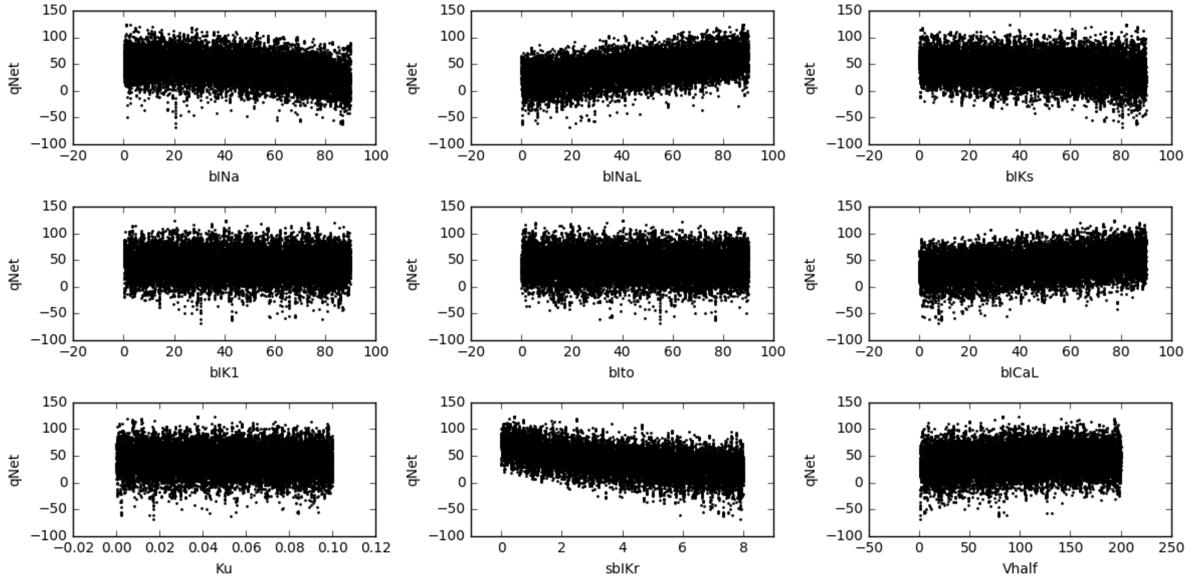


**Figure S1.** Scatter plot of $qNet$ verus different direct features for the 22000 simulated virtual drugs (endo cell model)

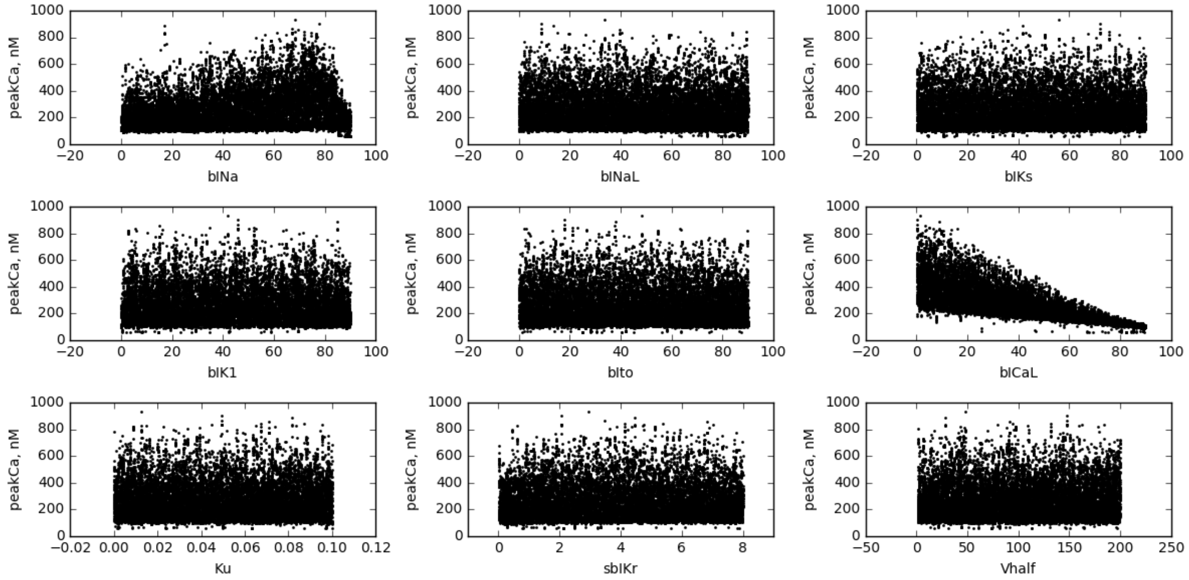


**Figure S2.** Scatter plot of $peakCa$ versus different direct features for the 22000 simulated virtual drugs (endo cell model)

#### Sobol sensitivity indices

The manuscript reports first-order ($S1$) and total sensitivity indices ($ST$) for the $qNet$, $APD90$ and $peakCa$ output metrics extracted from the endo cell model. Figures S3, S4, and S5 show the estimated values of the Sobol indices ($S1$ and $ST$) for eight derived features extracted from the endo, M, and epi cell type models, respectively.

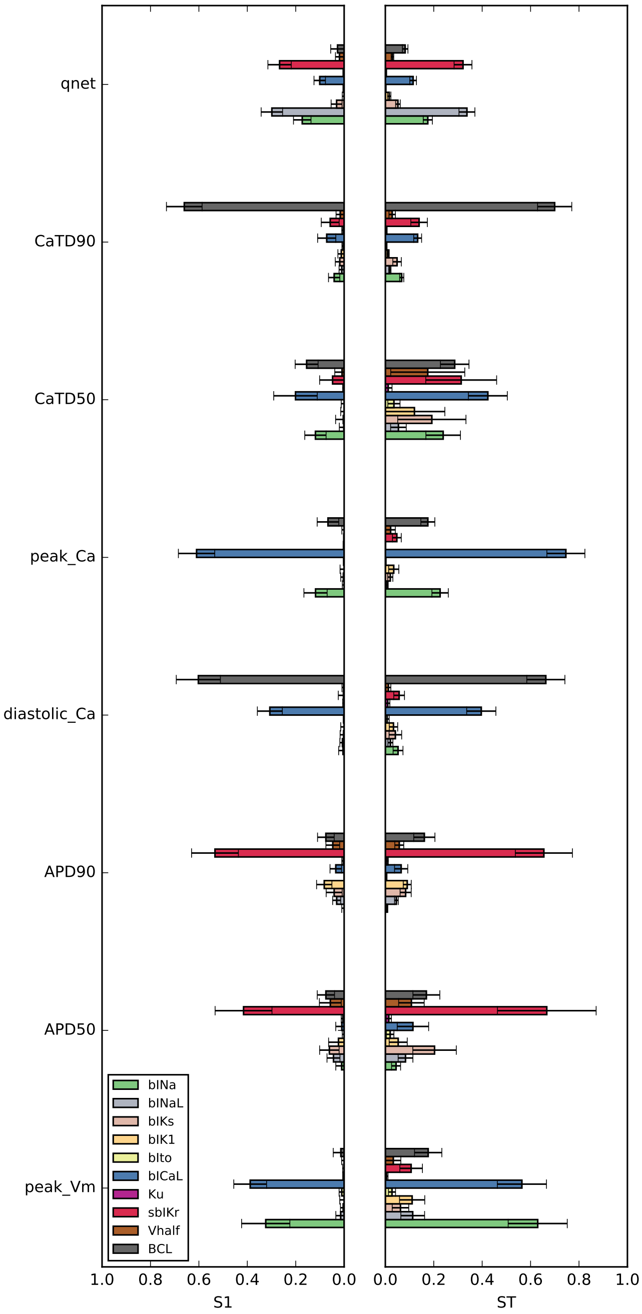


**Figure S3.** Plot of estimated first-order and total Sobol sensitivity indices (S1 and ST) to evaluate relative contributions of different input parameters on output variablity of 8 model-derived metrics (endo cell model)

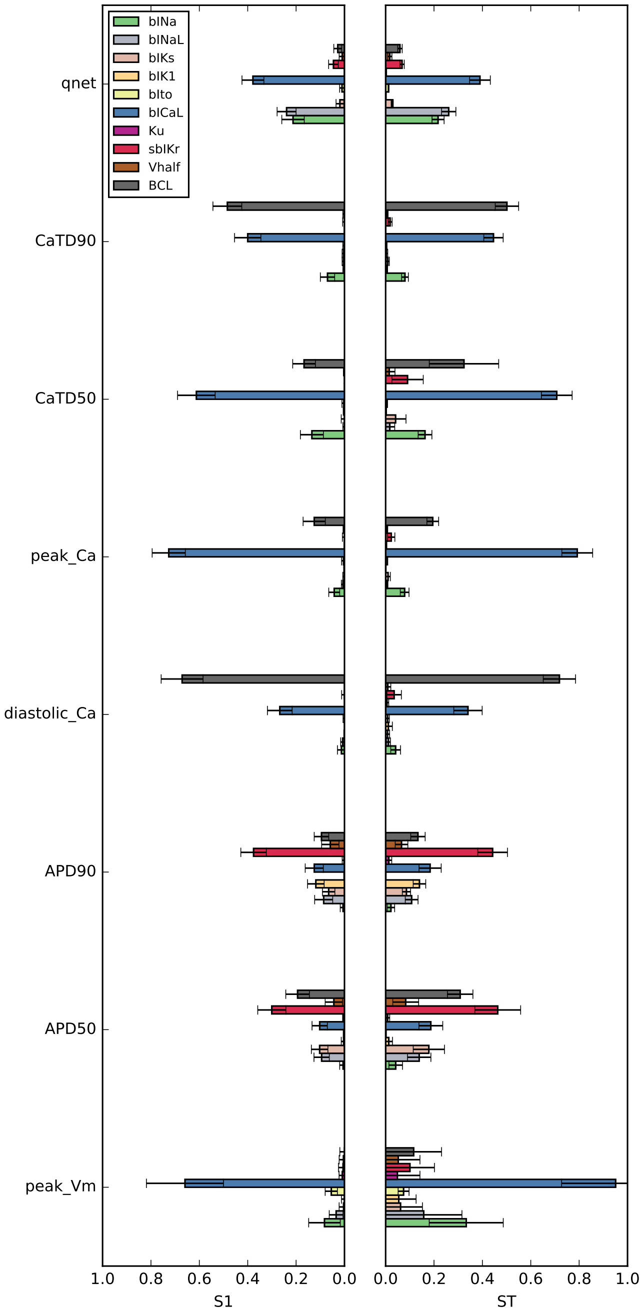


**Figure S4.** Plot of estimated first-order and total Sobol sensitivity indices (S1 and ST) to evaluate relative contribution of different input parameters on output variablity of 8 model-derived metrics (M cell model)

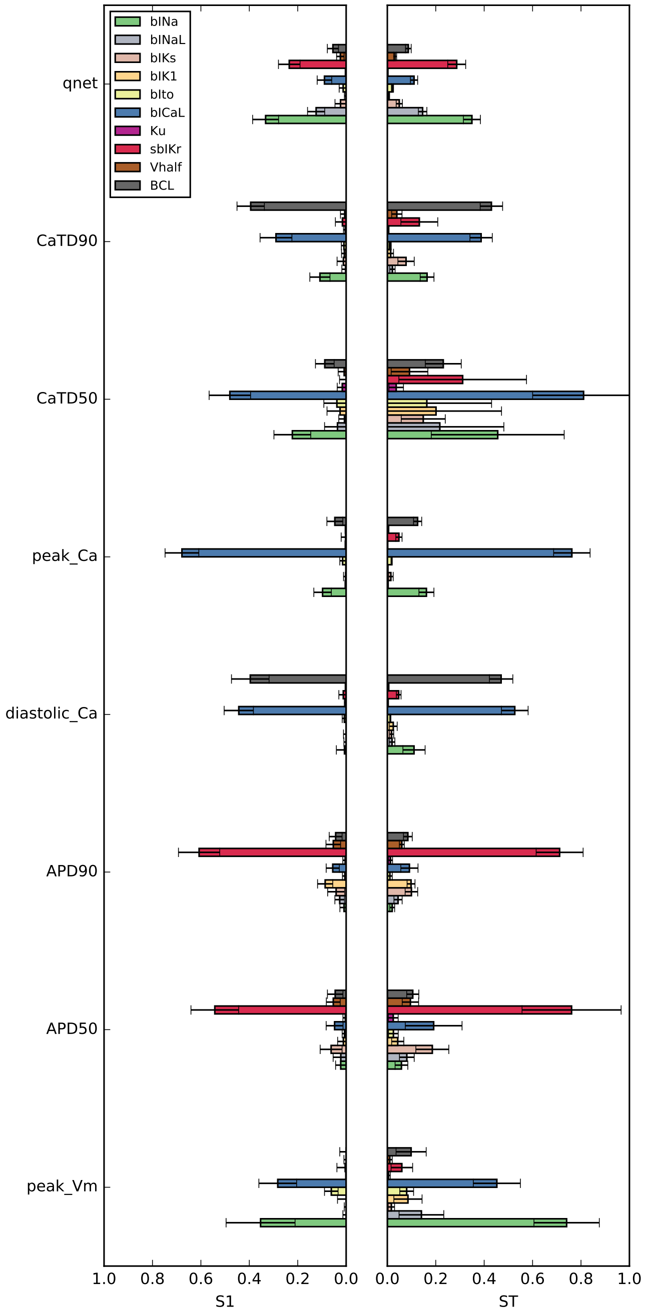


**Figure S5.** Plot of estimated first-order and total Sobol sensitivity indices (S1 and ST) to evaluate relative contribution of different input parameters on output variablity of 8 model-derived metrics (epi cell model)

#### Morris method

The Morris-method (Morris, 1991) provides the simplest form of GSA and can be considered as an extension of the local sensitivity methods. Similar to other local sensitivity analysis methods, the input parameters are varied one at a time (OAT) to obtain the partial derivatives of output metrics with respect to individual input parameters. A $k$-dimensional unit cube (representing $k$ independent normalized input parameters) is discretized into the $p$-level grid $\Omega$. Unlike for local sensitivity analysis, however, the reference values of each parameter are selected randomly from the set $\{0, 1/(p\text{-}1), 2/(p\text{-}1) \ldots , 1 - \Delta\}$ rather than perturbing some nominal values. The fixed increment $\Delta = p/2(p-1)$ is added to each parameter one at a time in a random order to compute the elementary effect (EE) of the $i^{th}$ input parameter $X_i$,

$$EE_i = \frac{Y(X_1, X_2, \cdots, X_{i-1}, X_i + \Delta, \cdots, X_k) - Y(X_1, X_2, \cdots, X_{i-1}, X_i, \cdots, X_k)}{\Delta}. \qquad \text{(S1)}$$

$k+1$ simulations are required to compute $EE_i$ for $k$ parameters. Multiple runs (paths) are performed from different start points to cover the entire parameteric space and obtain ensemble of EEs for each input parameter. The total number of simulations is $r \times (k+1)$, where $r$ is the number of paths. The mean of absolute $EE_i$ (denoted as $\mu^*$) and standard deviation of $EE_i$ (denoted as $\sigma_{EE}$) are computed for each input parameter. The $\mu^*$ statistic allows to rank the input parameters based on their influence (Saltelli et al., 2008) and $\sigma_{EE}$ helps identify nonlinear and/or interaction effects. The Python SALib package was used to perform the elementary effect analysis (Herman and Usher, 2017). In this study, we considered $r$ = 1000 unless otherwise specified, resulting in 9000 and 11000 input samples (virtual drugs) for $k$ = 8 and 10, respectively.

#### Morris method indices

#### Analysis of elementary effects

First, GSA was performed using the Morris method (see section Morris method) with $r = 1000$ repetitions and $k = 10$ input parameters, which require $n = r(k+1) = 11000$ model runs per cell type. Figure S6 shows relative impact of different input parameters on the variations of the $qNet$, $APD90$, and $peakCa$ output metrics for the endo cell. The X-axis represents $\mu^*_j$, a measure of the influence of the $j^{th}$ input variable on the output response. Larger $\mu^*$ values indicate greater influence of the input parameter on the output response. The Y-axis represents the standard deviation of the elementary effect $\sigma_j$. Higher $\sigma$ values for an input parameter indicate the presence of possible interactions with other variables affecting the output. The condition $\sigma/\mu^* > 1$ is an evidence of strong non-linear effects due to interactions between parameters. The input parameters with $\sigma/\mu^* < 0.1$ are instead considered to have minimal interactions. The analysis of elementary effects reveals that $bINaL$, $sbIKr$, $bINa$, and $bICaL$ have the strongest influence on the $qNet$ metric for the endo cell. Moreover, the ratio $\sigma/\mu^* < 1$ for these input parameters suggests that although they are not linearly correlated with $qNet$, they show a monotonic effect. On the contrary, the most influential input parameters affecting $APD90$ in the model was found to be $sbIKr$, the static component of the block of hERG channel. $peakCa$ concentration was most sensitive to the $bICaL$ parameter that represents the block of the L-type calcium channel current. The results suggest that blocks of different cardiac ion-channels (input parameters) have distinct influence on the derived metrics $APD90$, $qNet$, and $peakCa$ in the CiPAORd model.

The Figure S7 shows a plot of the $ST$ index against the $\mu^*$ statistic from the Morris method. The observed strong correlation is in agreement with previous analyses comparing elementary effects and variance-based methods (Campolongo et al., 2007). Given the results, the Morris method was able to rank the relative

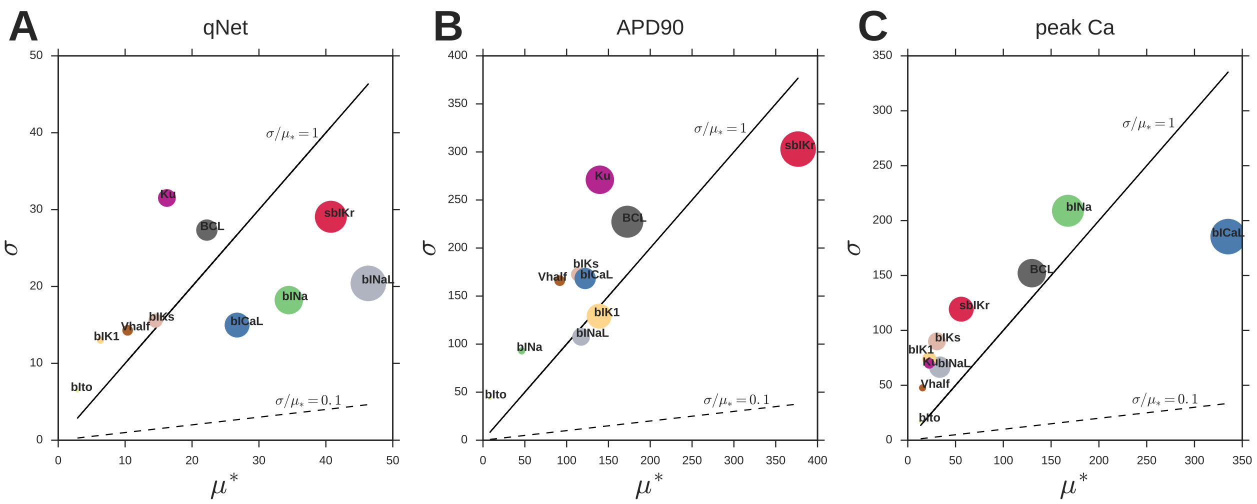


**Figure S6.** Estimators $\sigma$ and $\mu^*$ providing information on influence of blocks of different cardiac ion-channels, drug-binding parameters and BCL on CiPAORd endo cell model-derived metrics. **A**: $qNet$, **B**: $APD90$, and **C**: $peakCa$. The circle size for the input parameters correspond to their relative importance, with the largest diameter circle denoting the most influential input and the smallest diameter circle denoting the least influential inputs.

influence of parameters with similar accuracy of the more computationally expensive variance-based method. However, as discussed earlier, the variance-based sensitivity measures provide also additional information on parameter interactions.

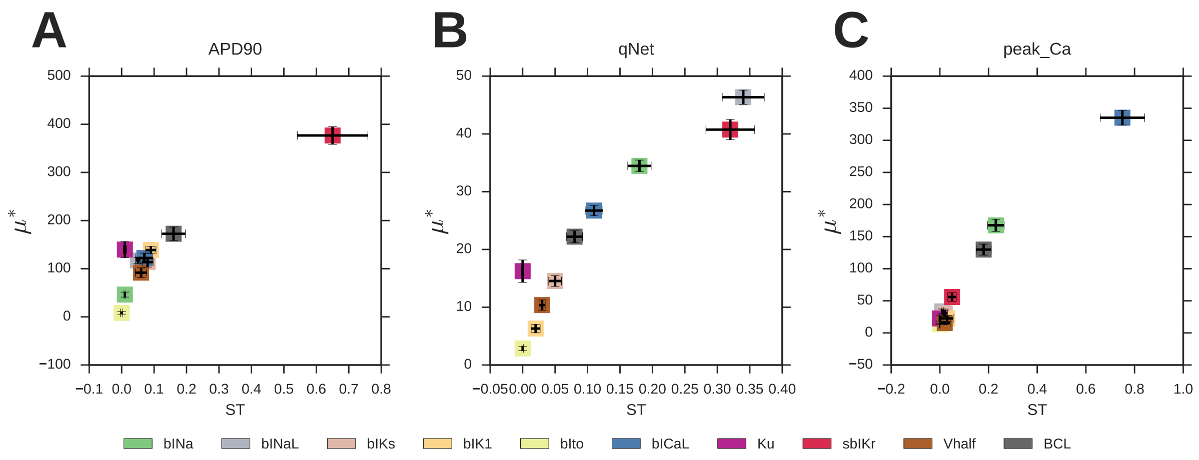


**Figure S7.** Plot of $\mu^*$ index calculated by Morris method versus $ST$ estimated in the Sobol sensitivity analysis. **A**: $qNet$ , **B**: $APD90$, and **C**: $peakCa$.

The estimated elementary effect values for all the derived features are shown in figures below.

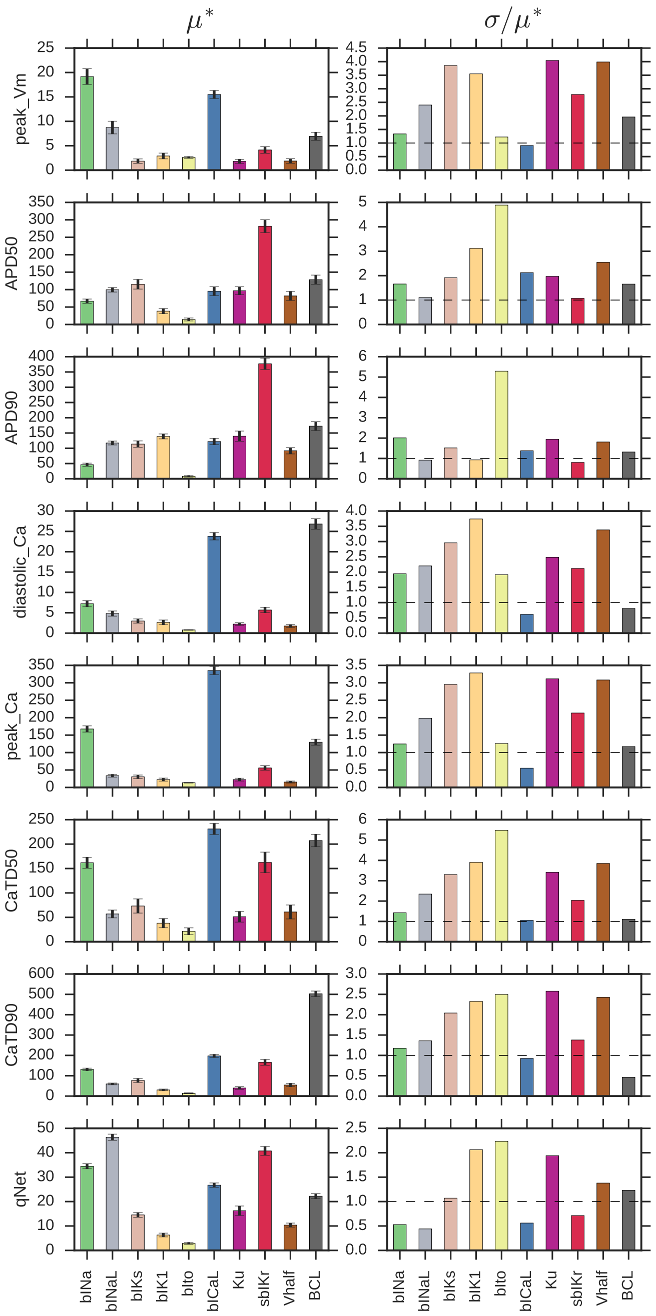


**Figure S8.** Plot of estimators $\mu*$ and $\sigma$, providing information on influence of input parameters on eight model-derived features (endo cell model)

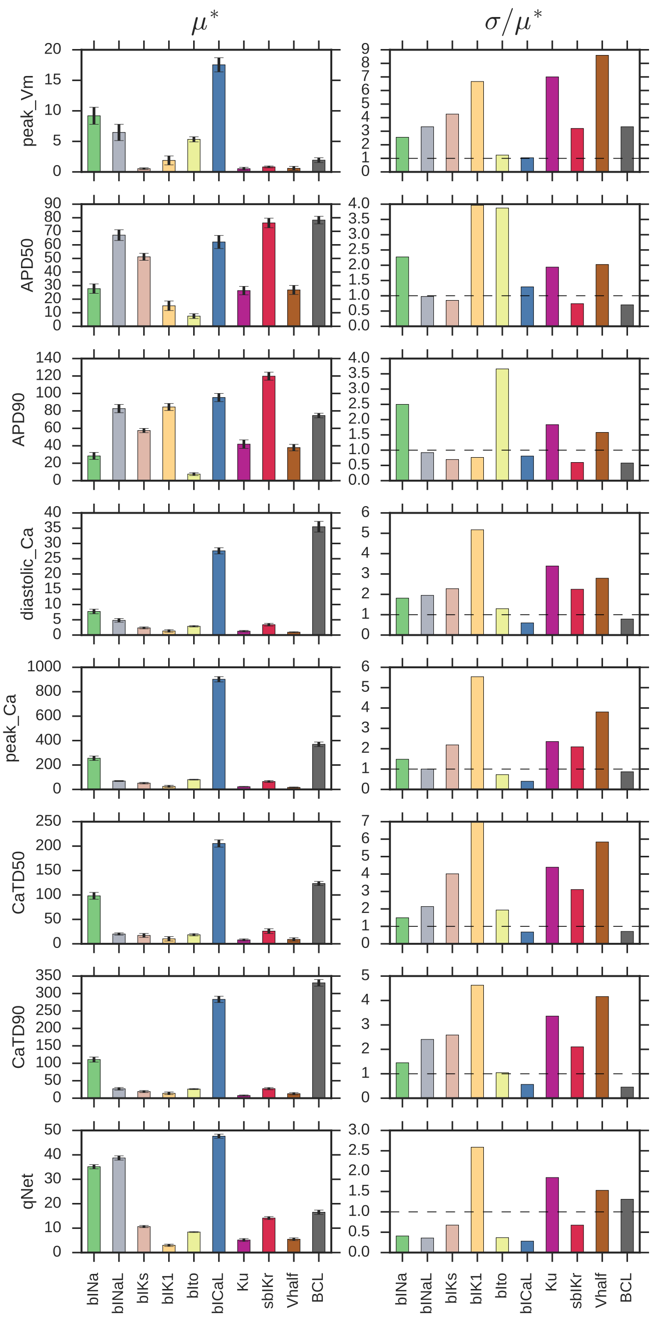


**Figure S9.** Plot of estimators $\mu*$ and $\sigma$, providing information on influence of input parameters on eight model-derived features (M cell model)

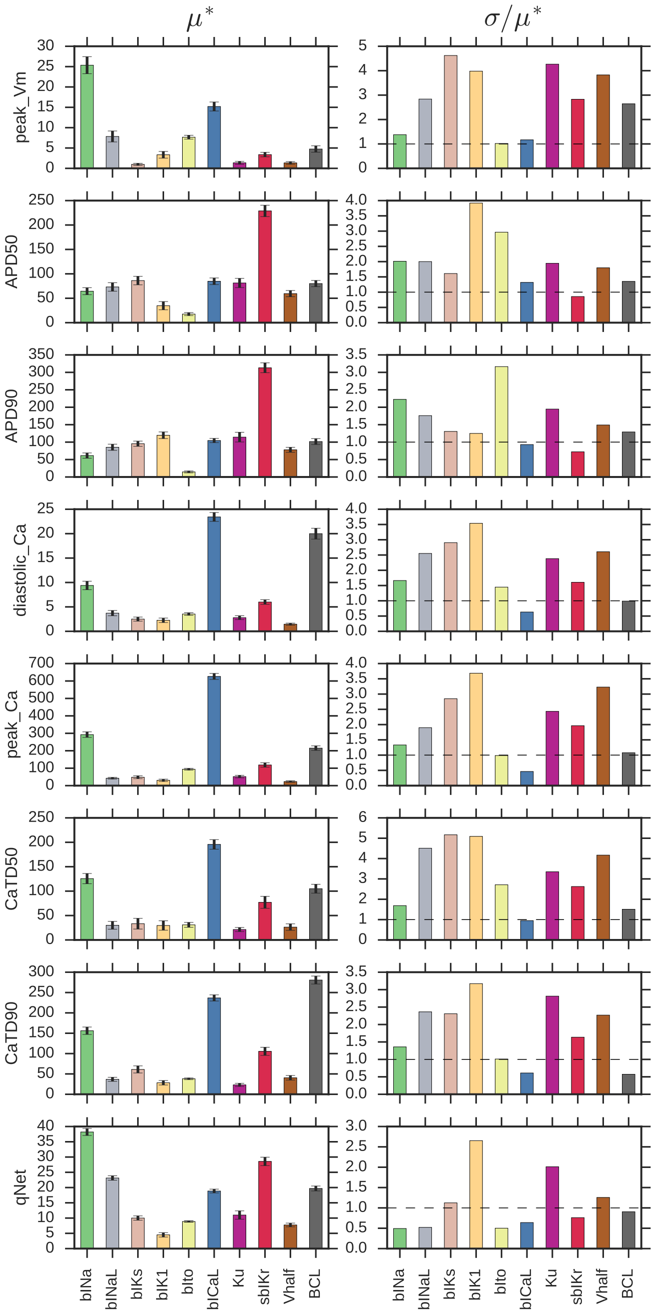


**Figure S10.** Plot of estimators $\mu*$ and $\sigma$, providing information on influence of input parameters on eight model-derived features (epi cell model)

#### Sobol sensitivity indices vs. sensitivity indices obtained from multivariate linear regression

The link between GSA and feature selection is well illustrated in the following example. We consider that some model-derived metric $M$ can be well approximated by a multivariatie quadratic polynomial, with direct features as input variables of the polynomial. The equation of the polynomial is

$$M = c + \sum_i a_i x_i + \sum_{ij} b_{ij} x_i x_j, \tag{S2}$$

where $\{x_i\}$ is a vector of direct features; $\{a_i\}$ and $\{b_{ij}\}$ are a vector and a symmetric matrix of coefficients for the linear and quadratic terms of the poylnomial, respectively, and $c$ is a constant. The components of $\{x_i\}$, i.e., the individual direct features, are assumed to be uniformly distributed with expectation $e_i$ and variance $v_i$. In this case, the effect of each direct feature $x_i$ and the joint effect of direct features $x_i x_j$ on variance in $M$ are given by

$$V_i = (a_i + \sum_{\substack{k \\ k \neq i}} 2b_{ik} e_i)^2 v_i + b_{ii}^2 var(x_i^2) + 2b_{ii}(a_i + \sum_{\substack{k \\ k \neq i}} 2b_{ik} e_i) cov(x_i, x_i^2), \tag{S3}$$

$$V_{ij} = 4b_{ij}^2 \left( v_i v_j + e_j^2 v_i + v_j e_i^2 - cov(x_i x_j, x_i) e_j - cov(x_j x_i, x_j) e_i \right). \tag{S4}$$

Typically, to rank the features based on their importance, feature scaling is carried out on the inputs before performing the regression. If each of the uniformly distributed direct features $x_i$ are centered and normalized to have $e_i = 0$ and $v_i = 1$, then

$$V_i = a_i^2 + \frac{4}{5} b_{ii}^2, \tag{S5}$$

$$V_{ij} = 4b_{ij}^2. \tag{S6}$$

Hence, the first- and second-order Sobol indices can be represented as

$$S1_i = \frac{a_i^2 + \frac{4}{5} b_{ii}^2}{var(M)}, \tag{S7}$$

$$S2_{ij} = \frac{4b_{ij}^2}{var(M)}. \tag{S8}$$

Expressions in (S7)-(S8) provide a link between calculated variances in the Sobol sensitivity method and the coefficients of the polynomial. Indeed, to demonstrate these relationships, we fitted a linear regression to approximate model-derived metrics ($APD90$, $qNet$, and $peakCa$) accounting for only the first-order effects of the direct features, i.e., $M = c + \sum a_i x_i$. Then, the squared coefficients of the regression are normalized by the variance of derived features and compared with Sobol $S1$ index. As expected, in the presence of minor interactions and nonlinear effects for these derived metrics, we observe a good match between indices estimated from the regression coefficients and the Sobol $S1$ indices as shown in Figure S11, even when considering only the first-order effects to construct the regression.

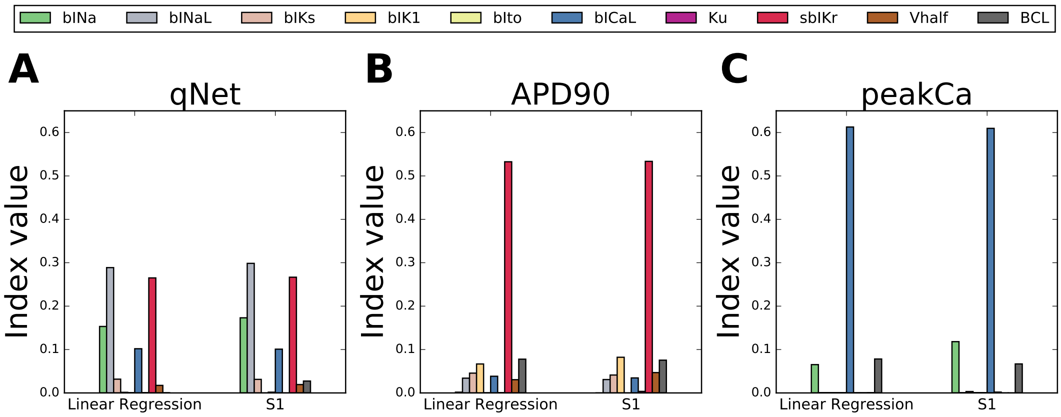


**Figure S11.** Comparision of index estimated from linear regression coefficients and Sobol first-order index $S1$ for **A:** $qNet$, **B:** $APD90$, and **C:** $peakCa$.

#### Simple examples highlighting differences between Mote Carlo Filtering versus Logistic Regression

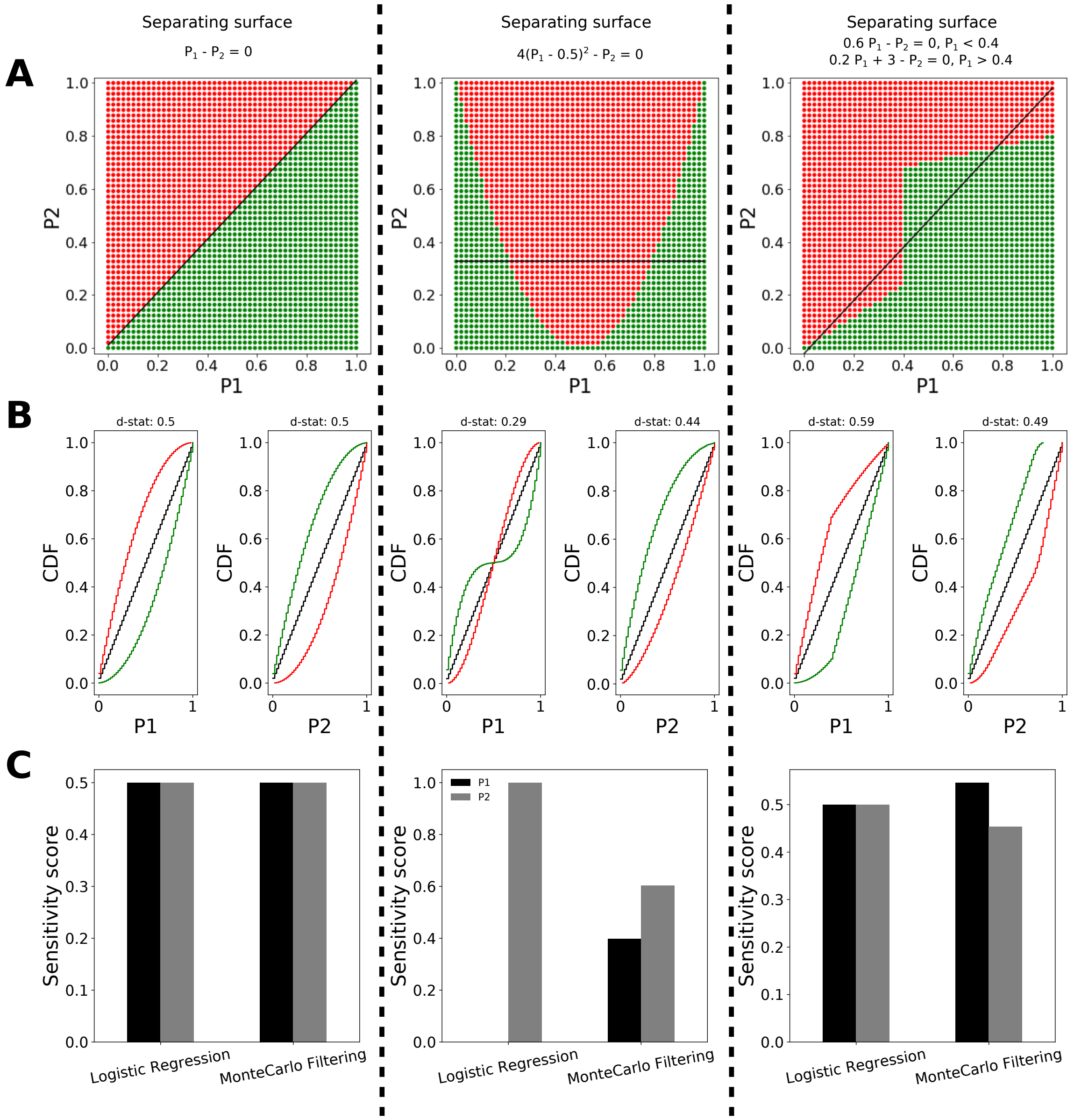


**Figure S12.** Comparision of sensitivity measures obtained via Monte Carlo Filtering and Logistic Regression Methods for hypothetical surfaces separating the behavioral and non-behavioral regions. **A:** Plots of three hypothetical separating surfaces. **B:** Empirical CDFs for both the input parameters $P1$ and $P2$. **C:** Bar plot comparing the obtained sensitivity indices with both the logistic regression and Monte Carlo filtering methods.

#### Comparision of EAD sensitivity based on Monte Carlo Filtering and Logistic Regression Methods

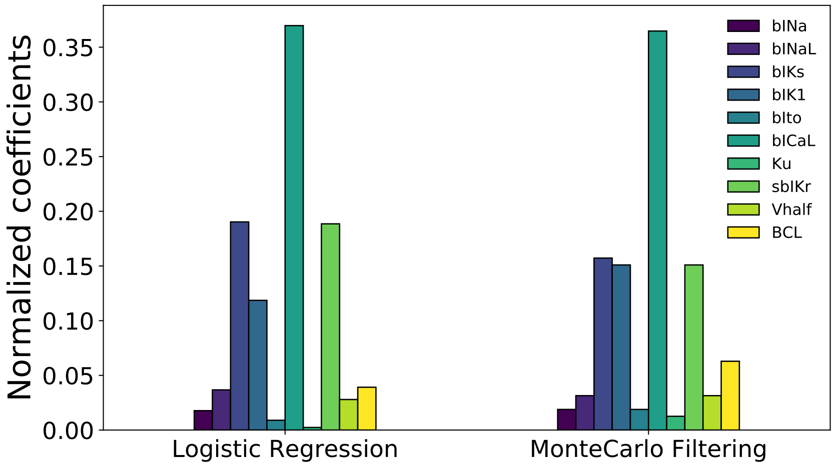


**Figure S13.** Comparision of obtained EAD sensitivity based on Montecarlo Filtering and Logistic Regression methods in the Endo cell CiPAORd model.

#### TdP risk classification of "CiPA drugs" based on transmural dispersion of repolarization

The "CiPA drugs" were also calssified based on the transmural dispersion of repolarizations as shown in Figure S14.

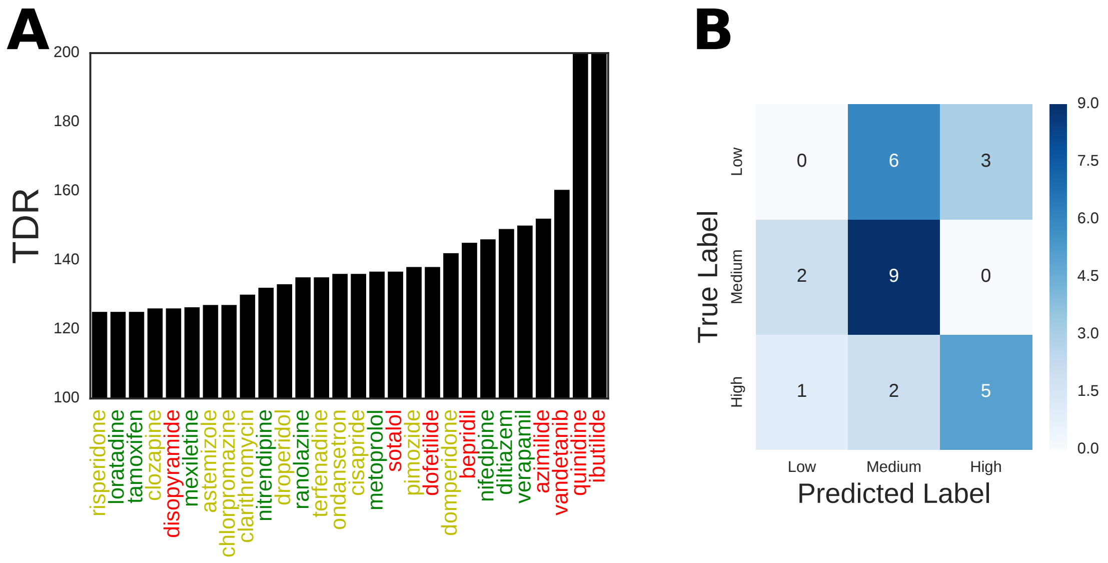


**Figure S14.** Risk categorization of "CiPA drugs" based on transmular dispersion of repolarization ($TDR$). **A:** $TDR$ values for the 28 CiPA training and validations compound at (2x Cmax drug concentrations) are estimated from simulations of the drugs at pacing rate of 2000 ms in the CiPAOrd model as difference between the $APD90$ values between the epi and the M cell type. **B:** Concordance analysis of $TDR$-based risk categorization of the 28 drugs against the actual torsadogenic risk of the compounds reported as confusion matrix.

#### Examination of pause-induced EADs in the ORd and CiPAORd models

A 3-D input parametric space was generated considering variations in the block of late sodium current ($INaL$), block of slow rectifitying potassium channel current ($IKs$) and L-type calcium channel currents ($ICaL$) at fixed value of static component of the hERG channel current ($sbIKr = 7$), as in our previous study (Parikh et al., 2017). Uniformly spaced samples in this 3D space were selected to generate a set of virtual drugs. The virtual drugs were simulated and outputs of the *in silico* models were visualized for presence or absence of EADs (Figure S15). The region in green in the Figure S15 denotes the region of the examined parameteric space where EADs were not observed in the CiPAOrd model, while the region in red denotes the region with presence of EADs in the model. The surface separating the $EAD+$ and $EAD-$ regions is shown in yellow for the CiPAORd model (Li et al., 2017). For comparision the estimated surface separating the $EAD+$ and $EAD-$ for the original ORd model (O'Hara et al., 2011) is shown in blue. The CiPA training compounds were plotted as red (High risk drugs), yellow (Intermediate risk drugs) and green (Low risk drugs) dots.

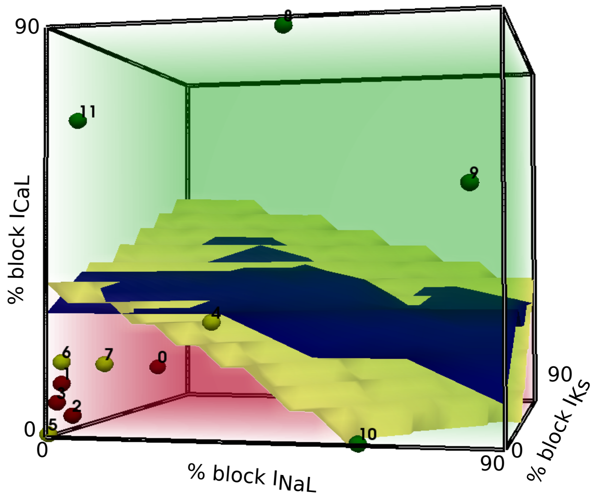


**Figure S15.** Surface separating the region of the input parametric space where EADs are observed in the simulated action potential from the region of the parameteric space where no EADs are observed. Blue surface: original ORd model and Yellow surface: CiPAORd model.
